# Dietary Restriction Impacts Peripheral Diurnal Rhythms Important for Longevity in *Drosophila*

**DOI:** 10.1101/2023.01.04.522718

**Authors:** Dae-Sung Hwangbo, Yong-Jae Kwon, Marta Iwanaszko, Shiju Sisobhan, Peng Jiang, Ladan Abbasi, Nicholas Wright, Sarayu Alli, Alan L. Hutchison, Aaron R. Dinner, Rosemary I Braun, Ravi Allada

**Affiliations:** Department of Neurobiology, Northwestern University, Evanston, IL 60208, USA; Biostatistics Division, Department of Preventive Medicine, Northwestern University, Chicago, IL 60611, USA; Michigan Neuroscience Institute, University of Michigan, Ann Arbor, MI 48109, USA; Center for Sleep & Circadian Biology, Department of Neurobiology, Northwestern University, Evanston, IL 60208, USA; James Franck Institute, Department of Chemistry, Institute for Biophysical Dynamics, University of Chicago, Chicago, IL 60637, USA; Department of Engineering Sciences and Applied Mathematics, Northwestern University, Evanston, IL 60208, USA; NSF-Simons Center for Quantitative Biology, Northwestern University, Evanston, IL 60208, USA; Department of Biology, University of Louisville, Louisville, KY 40292, USA

**Keywords:** Drosophila, Circadian Clock, Fat Body, Dietary Restriction, Aging, Proteasome, RNA-Seq

## Abstract

Circadian clocks may mediate lifespan extension by caloric or dietary restriction (DR). We find that the core clock transcription factor *Clock* is crucial for a robust longevity and fecundity response to DR in *Drosophila*. To identify clock-controlled mediators, we performed RNA-sequencing from abdominal fat bodies across the 24 h day after just 5 days under control or DR diets. In contrast to more chronic DR regimens, we did not detect significant changes in the rhythmic expression of core clock genes. Yet we discovered that DR induced de novo daily rhythmicity or increased expression of oscillating genes. Network analysis revealed that DR increased network connectivity in one module comprised of genes encoding proteasome subunits. Adult, fat body specific RNAi knockdown demonstrated that proteasome subunits contribute to DR-mediated lifespan extension. Thus, daily gene regulation links DR-mediated changes in rhythmic transcription to lifespan extension.

## Introduction

Circadian (∼24 h) clocks regulate a wide range of rhythmic metabolic, physiological and behavioral parameters to acclimate to environmental changes in light, temperature, and food availability (Patke et al. 2020). Circadian clock disruption has been implicated in advanced aging and the longevity response to caloric or dietary restriction (CR or DR) (Galikova and Flatt 2010; Manoogian and Panda 2017; Froy 2018; Nakahata and Fukada 2022; Zhu et al. 2022). DR, reduction in food intake without causing malnutrition, robustly extends longevity in various animal models including yeast, worms, flies, and monkeys (Green et al. 2022; Mc Auley 2022). Yet, the molecular mechanisms by which DR delays aging are not fully understood. Understanding how the clock impacts aging and DR sensitivity may provide novel avenues to understanding aging.

The circadian clock consists of a widely conserved transcriptional feedback loop that drives 24 h molecular oscillations. In flies, the heterodimer transcription factor CLK/CYC forms the positive arm of the loop and activates their repressors, PER and TIM. The PER-TIM complex functions as the negative arm of the loop and inhibits CLK-CYC activity (Allada and Chung 2010). This feedback loop drives core clock gene rhythms and controls rhythmic physiological, metabolic, and behavioral parameters via clock control of output genes (Patke et al. 2020). Genetically hybrid mice with a deviation of the circadian period from 24 h by over seven minutes showed a higher mortality rate than the mice with less deviated periods (Libert et al. 2012). However, whether the altered circadian period is correlated with or causes the increased mortality is not clear. Genetic inactivation of CYC ortholog *Bmal1* as well as other circadian clock mutants also significantly reduced lifespan in mice (Fu et al. 2002; Kondratov et al. 2006; Dubrovsky et al. 2010; Lee et al. 2010). Yet when *Bmal1* knockout was restricted to adulthood, lifespan was normal (Yang et al. 2016). While a lifelong DR did not significantly extend lifespan of *Bmal1* knockout mice (Patel et al. 2016a), chronic (∼2 mo) DR exposure increased core clock amplitude in mice (Patel et al. 2016b; Sato et al. 2017). This suggests that the circadian clock may be among the molecular mechanisms of DR. However, mice under DR restrict their feeding behavior to a narrow temporal window (Acosta-Rodriguez et al. 2017). Thus, DR induced changes in core clocks may instead be due to the well-known effects of time-restricted feeding (Hatori et al. 2012). Indeed, core clock genes are important for age-dependent cardiac function and lifespan extending effects of time-restricted feeding (Gill et al. 2015; Ulgherait et al. 2021). A study revealed that a basal level lifespan extension by DR is further increased when DR is temporally aligned with mice’s natural meal timing (i.e., during the night) (Acosta-Rodriguez et al. 2022). Thus, it remains unclear whether disruption of the circadian clock itself or other factors, such as a defect during development, results in lifespan reduction and is responsible for the lack of DR response.

Circadian clocks have also been implicated in aging and the DR longevity response in flies as well. DR mortality effects are rapid, fully evident within just 2 ∼ 4 days of a diet shift in flies (Mair et al. 2003; McCracken et al. 2020), making them an attractive model organism for DR studies. Loss-of-function mutants in the activator and repressor complexes that “fix” the clock at different points in the cycle have tested the functional significance of the clock in aging and DR. Inhibition of neuronal *Clk* appears to reduce the lifespan extending DR effects, where flies were tested for DR effects with just two (ad libitum and DR) diets (Hodge et al. 2022). However, this observation is inconclusive to the role of *Clk* for DR effects as inhibition of neuronal *Clk* decreases food intake (Xu et al. 2008). Reduction of food intake can decrease lifespan under DR while increasing lifespan under ad libitum, masking the true DR response (Flatt 2014). *per^01^*and *tim^01^* mutant flies exhibit inconsistent DR longevity responses perhaps due to differences in microbial content (Katewa et al. 2016; Ulgherait et al. 2016; Ulgherait et al. 2020; Ulgherait et al. 2021). Thus, the role of core clock genes in mediating DR effects could be further clarified.

Notably, the rhythmic amplitude of core clock genes of flies is enhanced after chronic (>10 days) DR (Katewa et al. 2016). Knockdown of modestly CLK- and DR-regulated genes in the eye modulate lifespan without apparent effects on DR-dependent longevity (Hodge et al. 2022). *tim* overexpression increased the amplitude of core clock oscillations and extended lifespan under control but not DR diets (Katewa et al. 2016). While clock oscillation amplitudes between *tim* overexpression on a control diet and wild-type flies under DR are comparable, their lifespan remain quite different, suggesting that core clock effects may not be required for lifespan extension (Katewa et al. 2016). Thus, it remains unclear if lifespan extension functions via the circadian clock or instead through clock output genes and what the role of specific clock output genes is in DR-dependent longevity.

Using multiple diets, we demonstrate that *Clk* mutants suppress DR longevity and fecundity responses, providing more definitive demonstration of the role of the core clock. Nonetheless, using a shorter-term DR strategy, we reveal that primary DR effects on the circadian transcriptome spare core clock genes, suggesting a primary effect on rhythmic output. Network analysis suggests that a diet-dependent effect on a gene module containing proteasome subunit genes. Moreover, suppression of proteasome subunit expression, predominantly in the abdominal fat body, limits lifespan extension by DR. These results provide crucial genetic evidence that circadian clock output pathways, specifically those involving the proteasome, link DR-mediated changes in rhythmic transcription to lifespan extension. These studies raise the possibility of using chronotherapy to combat aging and age-related diseases.

## Results

### The Effects of Dietary Restriction on Lifespan and Fecundity Is Dramatically Suppressed in Mutants of the Core Clock Transcription Factor *Clk*

To tease apart the role of the circadian clock in the DR longevity response, we first evaluated the roles of the positive and negative arm of the feedback loop by testing *per^01^* as in prior reports (Katewa et al. 2016; Ulgherait et al. 2016; Ulgherait et al. 2020), and *Clk^Jrk^*, a dominant negative allele of *Clk* (Allada et al. 1998), which had not been previously examined for DR studies. The *Clk^Jrk^*allele encodes a premature stop codon removing a putative transcriptional activation domain but preserving its DNA-binding and dimerization motifs (Allada et al. 1998). Thus, *Clk^Jrk^* is predicted to behave as a dominant negative minimizing compensatory effects as observed for pure loss-of-function *Clock* mutants in mice (Debruyne et al. 2006). We applied a common DR regimen where the concentration of both yeast and sucrose are diluted (whole food dilution: Control: 15% [w/v] Sucrose and Yeast, 15SY; DR: 5% [w/v] Sucrose and Yeast, 5SY) in 12hr light: 12hr dark (LD) cycles (Bass et al. 2007; Kabil et al. 2011). We used female flies where DR responses are more robust (Magwere et al. 2004). We observed robust lifespan extension by DR in wild-type *iso31 (w^1118^,* 29%) and comparable extension in *per^01^*flies (25%) (Figure 1-figure supplement 1; diet× genotype interaction p > 0.05 between *iso31* and *per^01^*). Here we use the term wild-type as wild-type for the locus of interest in comparison to a genetic mutant or perturbation. These results are consistent with one report which showed that *per^01^*flies display normal DR extension effects (Ulgherait et al. 2016). However, only ∼ 10% lifespan extension by DR was observed in *Clk^Jrk^* mutants (Figure 1-figure supplement 1; diet×genotype interaction p < 0.0001 between *iso3*1 and *Clk^Jrk^*). Interestingly, the DR responses in *per^01^* and *Clk^Jrk^* flies were significantly different from each other (Figure 1-figure supplement 1; diet×genotype interaction p < 0.0001 between *per^01^* and *Clk^Jrk^*). We hypothesize that *Clk^Jrk^* and *per^01^* arrest the clock at opposite points in the cycle, only one of which impacts the DR response.

**Figure 1.**
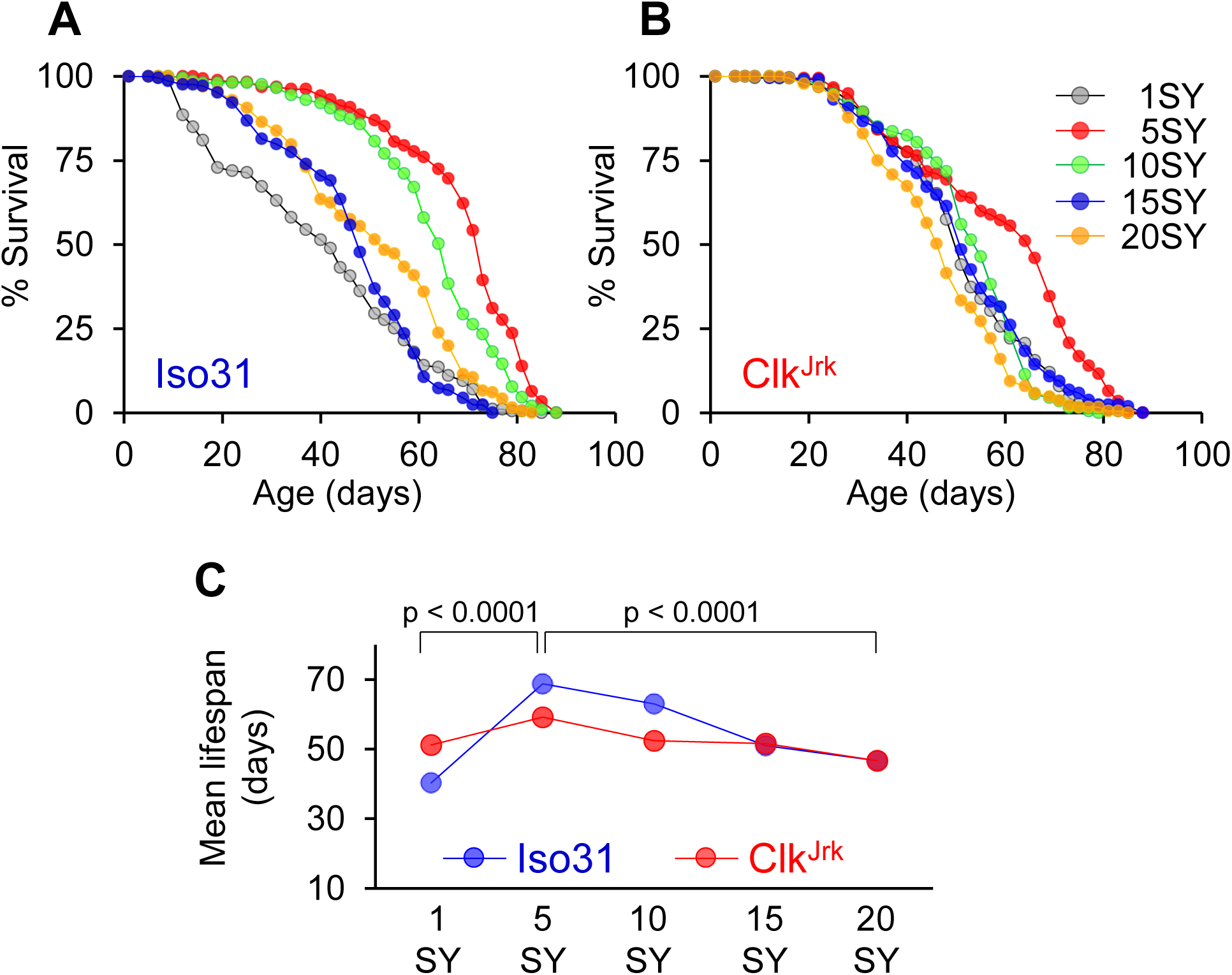
**Reduced longevity response to DR in *Clk^Jrk^* mutant flies** (A-B) Survival curves of wild-type control flies (*iso31*) and *Clk^Jrk^*homozygous mutant flies in 1%, 5%, 10%, 15%, and 20% sucrose-yeast (SY) diets (total dilution). (C) Mean lifespan plots of *iso31* and *Clk^Jrk^* flies across different concentrations of SY diets. Basic survival parameters from the Kaplan-Meier method for each diet and genotype are in Supplementary File 1. P values represent the probability of main (Gene (G) as a nominal variable and Diet (D) as a continuous variable) and interaction effects (G x D) by the likelihood ratio chi-square test from Cox proportional hazards regression analysis to evaluate ability of tested genes to modify lifespan in the specified range of diets. Separate analyses were performed for normal DR response (from 5SY to 20SY) and malnutrition-like response (from 5SY to 1SY). An independent replication of the experiment is presented in Figure 1-figure supplement 2. See Supplementary File 1 for additional details of the statistical analysis.

Although the DR response is typically tested by comparing lifespan with just two diets (Solovev et al. 2019), it can potentially mis-assign the effects of DR or diet (Tatar 2007; Flatt 2014). For example, *chico* mutants show little effect of DR looking at just two diet concentrations but showed a robust DR response if their lifespan was measured across seven diet concentrations (Clancy et al. 2002). To distinguish between this possibility and a “true” DR response, we performed a reaction norm analysis by comparing the mean lifespan of *Clk^Jrk^* flies to that of wild-type over serially diluted diets (1, 5, 10, 15, and 20SY) (Bass et al. 2007; Tatar 2007; Flatt 2014). As expected in wild-type flies, we observed a reduction of lifespan as food concentration increased from 5SY to 20SY. A reduction of lifespan was also observed when going from 5SY to 1SY, presumably due to malnutrition (Figure 1A, C). However, mean lifespan of *Clk^Jrk^* mutant flies showed a more flattened reaction curve to diets (Figure 1C, diet×genotype interaction p < 0.0001 between wild-type *iso31* and *Clk^Jrk^*), suggesting that *Clk* is a “true” DR gene. For example, *Clk^Jrk^* flies only show a 14% increase in lifespan between 5SY and 15SY while *iso31* flies show a 34% increase. In addition, while *Clk^Jrk^* flies are short-lived relative to wild-type at 5SY, they significantly outlived wild-type flies in 1SY (Figure 1, Figure 1-figure supplement 2, p < 0.0005 by log-rank test). Similar results were obtained in an independent trial (Figure 1-figure supplement 2).

To rule out the possibility that *Clk^Jrk^* effects could be explained compensatory feeding, we assessed food intake in ∼1 week old flies on three diets (1SY, 5SY, 15SY) over two days (48 hours) using the Con-Ex method (Shell et al. 2018). We observed either no significant changes (5SY and 15SY) or a relatively modest increase (1SY) in food consumption between *iso31* controls and *Clk^Jrk^* flies that are limited compared to the large differences in caloric content between the diets (Figure 1-figure supplement 3). While we cannot rule out changes in feeding patterns throughout the lifespan, we believe that minimal differential compensatory feeding in the *Clk^Jrk^* mutants is not sufficient to be the primary cause of the lifespan differences observed across diets.

It has been suggested that a whole food dilution may cause dehydration effects, especially on high concentrations of yeast (Ja et al. 2009) and sucrose (van Dam et al. 2020), complicating the interpretation of diet-lifespan effects. To address this possibility, we tested the DR response of *Clk^Jrk^* mutant flies using the yeast-restriction strategy, where varying yeast concentration with a fixed sucrose concentration (Bass et al. 2007; Ja et al. 2009; McCracken et al. 2020). We first tested the DR response of *Clk^Jrk^* mutant flies using the same experimental diets used for Figure 1 except with a fixed sucrose concentration at 5% (Figure 1-figure supplement 4). Although the reaction pattern of *Clk^Jrk^* flies in this yeast-restriction DR regimen was qualitatively somewhat different from that of whole food dilution, we confirmed that the DR response of the mutants was strongly suppressed as in whole food dilution protocols (Figure 1, Figure 1-figure supplement 2). We then further confirmed this observation using a different yeast-restriction protocol where purified yeast extract is used (Katewa et al. 2016; Ulgherait et al. 2016) instead of whole-cell lysates of yeast (Min et al. 2007; McCracken et al. 2020). Although overall mean lifespan and lifespan response to the yeast extract diets (Figure 1-figure supplement 5, Supplementary File 1) was qualitatively different from those of whole food restriction (Figure 1, Figure 1-figure supplement 2) and whole cell lysates yeast restriction (Figure 1-figure supplement 4), DR response of *Clk^Jrk^* mutant flies in yeast extract diets was significantly impaired compared to wild-type *iso31* control flies. Thus, across several diet conditions, these data indicate that *Clk^Jrk^*robustly suppresses the DR longevity response.

While increasing diet concentration has a negative impact on lifespan, it has a strong positive correlation with female fecundity (e.g., egg laying) primarily due to increased protein sources in yeast (Bass et al. 2007; Skorupa et al. 2008). To determine if *Clk^Jrk^* also impacted this diet response, we measured egg laying in both *iso31* and *Clk^Jrk^*mutant flies over 7 days. As expected, *iso31* flies increased egg production with increased diet concentrations (Figure 2). However, egg production was strongly decreased in *Clk^Jrk^* mutant flies (Figure 2, p < 0.0001 by regression analysis). More importantly, diet-dependent increases in egg laying were significantly suppressed in *Clk^Jrk^*mutants (Figure 2, p < 0.01 for pair-wise comparison in each diet by t-test). A similar trend was observed in an independent trial (Figure 2-figure supplement 1). Taken together, these data indicate that *Clk^Jrk^* strongly disrupts how flies respond to DR at both levels of longevity and fecundity.

**Figure 2.**
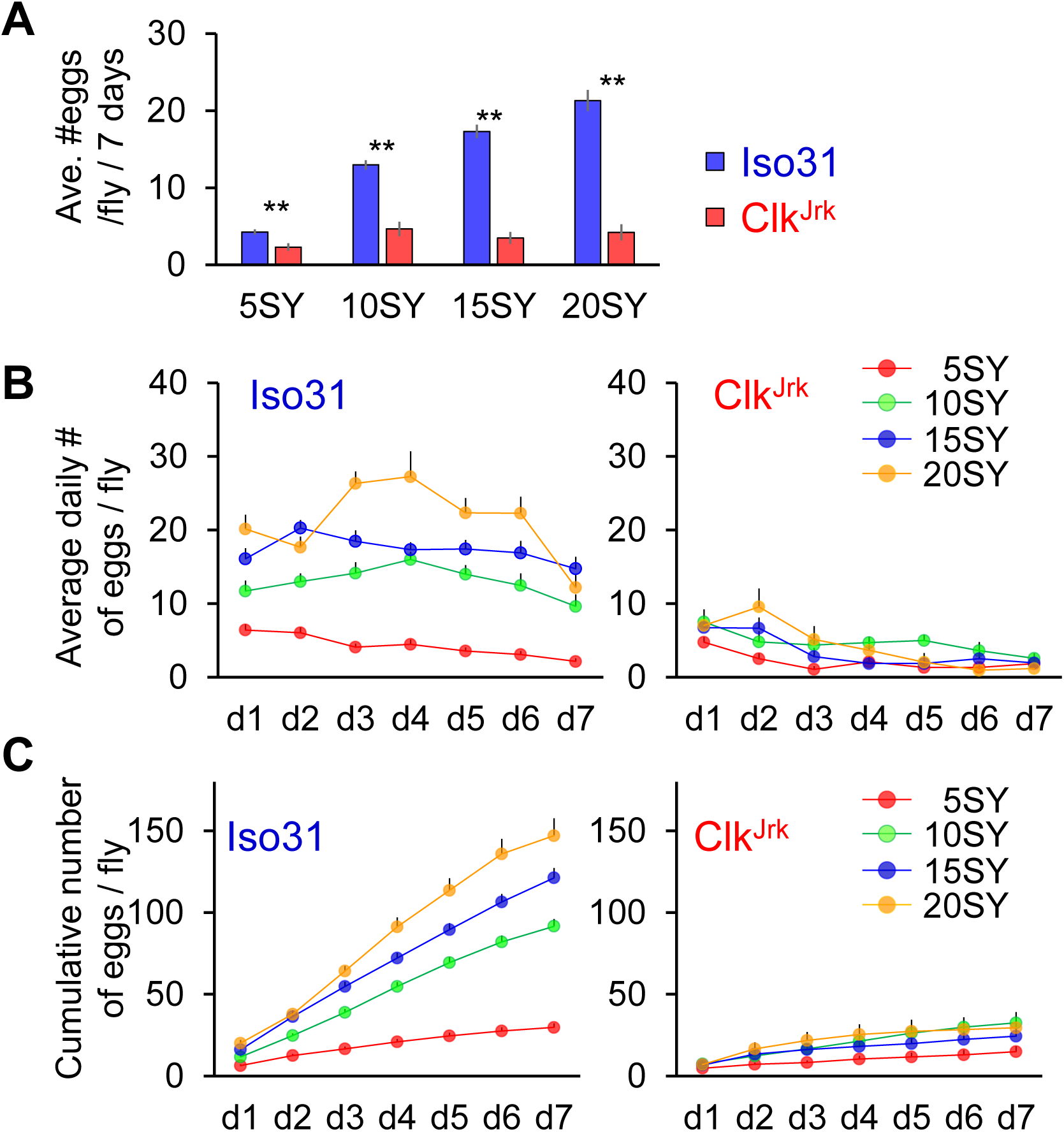
**Reduced fecundity response to diets in *Clk^Jrk^* mutant flies.** (A and B) Cumulative and daily average number of eggs produced per fly over 7 days in wild-type control flies (*iso31*) and *Clk^Jrk^* homozygous mutant flies on 5%, 10%, 15%, and 20% sucrose-yeast (SY) diets. (C) Cumulative number of eggs produced per fly over 7 days. ** p < 0.01 by t-test for specified pairwise comparisons. All error bars represent SEM.

### Shorter Term Dietary Restriction Selectively Reprograms Daily Rhythmic Genes in the Abdominal Fat Body

Given the role of *Clk^Jrk^* in mediating responses to DR, we then asked how the daily transcriptome responds to DR. RNA-Seq analysis using whole-fly lysates of female flies after seven days of DR or *ad libitum* diets (a yeast extract restriction protocol) suggested that DR alters the daily transcriptome in whole flies (Hodge et al. 2022). We focused our studies on the fat body, an analog of the mammalian liver and adipose tissue, given its critical role in mediating the effect of DR (Bai et al. 2012; Banerjee et al. 2012; Katewa et al. 2016; Dobson et al. 2018). Importantly, the fat body also has its own core clock system, including *Clk*, regulating energy metabolism, feeding, and egg-laying (Xu et al. 2008; Xu et al. 2011). To assess DR-dependent circadian rhythms, we performed a fine scale (every 2 h over 24 h) RNA-Seq analysis from dissected abdominal fat body tissues (Xu et al. 2008; DiAngelo and Birnbaum 2009; Xu et al. 2011) of young (8 days old) *iso31* females. To prepare dissected fat body samples for RNA-Seq analysis, we entrained flies for ∼ 5 days under 12h:12h light-dark (12LD) cycles on either control or DR diets. Previous studies showed that ∼ 3 days in LD cycles are sufficient to entrain the fat body clock in *Drosophila* (Xu et al. 2008; Xu et al. 2011; Erion et al. 2016). Moreover, DR reshapes mortality of flies within 2 ∼ 4 days of diet shift in flies (Mair et al. 2003; McCracken et al. 2020). We also observed that 5 days on DR diet was sufficient to affect metabolism and physiology evidenced by changes in fecundity (Figure 2, Figure 2-figure supplement 1). Thus, our environmental settings in light schedule and diet are sufficient to capture significant diet- and circadian-dependent transcriptional changes important for lifespan extension by DR.

We then analyzed the samples from control and DR diets separately to examine if and how DR changes the daily gene expression pattern in the fat body. For this diet-dependent analysis, we used RAIN (Thaben and Westermark 2014) to identify rhythmic genes, using a log2 fold change > 0.6 at an FDR <0.1. This cutoff corresponds to p-values of 0.003 and 0.005 for control and DR., making the p-value cutoff for this more stringent than those used in several studies (Eckel-Mahan et al. 2013; Abruzzi et al. 2017; Kuintzle et al. 2017; Sato et al. 2017). We found a significant reorganization of the diurnal transcriptome by DR (Figure 3B, C, E-G). Collectively, we identified 652 oscillating in either one or both conditions, of which 115 cycle in both conditions, 175 cycle only in the control diet, and 362 cycle only in the DR diet. Thus, there is a net increase of 64 % in cycling genes under DR (Figure 3B, C). As fat body samples were collected under a standard 12L:12D diurnal cycle, we compared them with *Clk^Jrk^* mutant flies to evaluate whether LD-rhythmic genes in wild-type flies are genuinely controlled by the circadian clock. Overall, we found under control conditions that the far majority of the cycling genes (279 out of 290 cycling genes) as well as the core clock genes are not detectably oscillating in *Clk^Jrk^*, suggesting that they are clock-driven (Figure 3-figure supplement 1).

**Figure 3.**
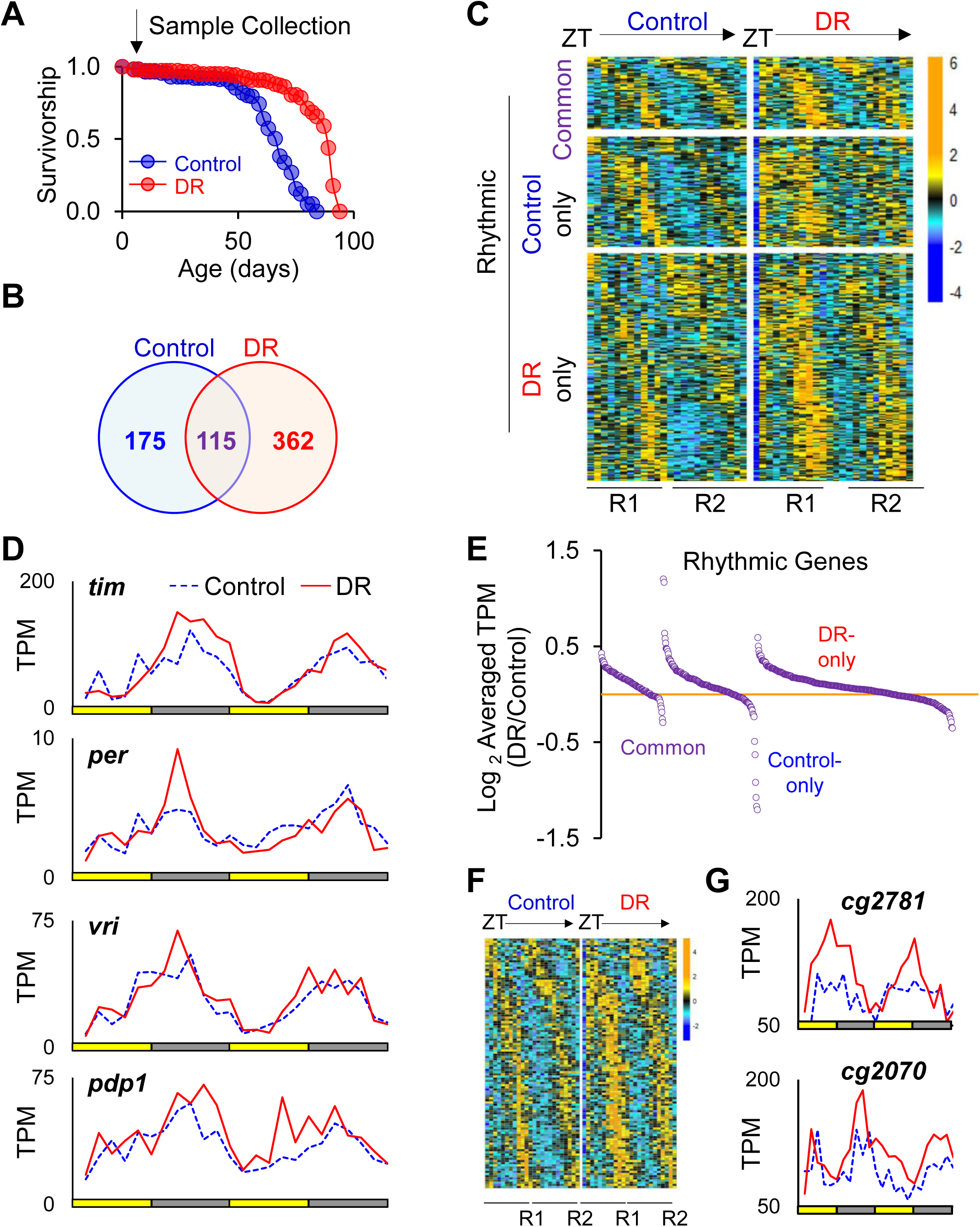
**Effects of DR on circadian transcriptome in the abdominal fat body** (A) Lifespan extension by DR. Samples for RNA-Seq analysis were collected after ∼5 days under either control or DR diets. (B) Number of rhythmic genes (RAIN, FDR<0.1 and log2 fold change >0.6). (C) Reorganization of the diurnal transcriptome by DR. The heatmap represents the relative expression (Z-score) of rhythmic genes in each group across 48 h at 2 h intervals (2 replicates of 12 time points over 24 h). Genes in the top panels are rhythmic in both control and DR diets (common); those in the middle panels are rhythmic in the control diet (left) but arrhythmic in the DR diet (right); those in the bottom panels are rhythmic in the DR diet (right) but arrhythmic in the control diet (left). (D) DR failed to affect the expression patterns of core clock genes. (E) Effect of DR on overall time-averaged expression of rhythmic genes in each group. TPMs of rhythmic genes in each group were averaged across all time points and normalized to those in the control diet. (F) Increased overall expression in the common rhythmic genes under DR. The heat map represents relative expression (Z-score) of common rhythmic genes across all time points from both control and DR diets. (G) Examples of common rhythmic genes with increased expression under DR.

Remarkably, core clock genes (e.g., *per*, *tim* and *Clk*) were not significantly impacted by DR in both phase and peak expression (using thresholds FDR < 0.1 and log2 (fold change) > 0.5 from differential analysis, see methods) (Figure 3D). This indicates that rhythmicity of core clock genes is largely resistant to short term (∼5 days) diet changes. Although peak phases of common rhythmic genes remained largely unaffected (Figure 3C), DR significantly, albeit modestly, increased overall expression of many of these common cyclers (Figure 3E-G). From an averaged expression comparison across all time points between control and DR in the 115 common rhythmic genes, 29 genes were significantly increased (paired t-test, FDR < 0.1) while none was downregulated (Figure 3E-G). It shows that short-term DR (5 days) increases expression of robustly rhythmic genes without affecting expression of the core clock genes, arguing that a short-term DR for ∼ 5 days impacts rhythmic output while sparing core clocks.

### Weighted Gene Coexpression Network Analysis Identifies a DR-Specific Cycling Proteasome Module

In order to identify novel genes and pathways that are associated with or even causal to the lifespan extension under DR and also to understand diurnal transcriptomic organization in the fat body under DR, we took a network approach (Zhang et al. 2013). First, we performed Weighted Gene Coexpression Network Analysis (WGCNA) to reconstruct gene coexpression networks from our time-series RNA-Seq data collected under DR diet (See Methods). We identified 46 network modules of co-regulated genes (Figure 4A, Supplementary File 2). Notably, 14 modules (30%) were “cycling modules”, i.e., those enriched with rhythmic genes (FDR < 0.05 and odd ratio > 0, Fisher’s exact test) (Figure 4B). Genes in each of these “cycling modules” exhibited highly similar phases and waveforms, revealing coordinated circadian gene expression (Figure 4C). In order to further examine how the transcriptomic organization is altered by DR, we computed the modular differential connectivity (MDC) (Zhang et al. 2013), which was expressed as a ratio to reflect the difference in gene co-expression strength (i.e., network connectivity) of a module between DR and control diets (see Methods and Supplementary Information). Among the 46 network modules, At FDR < 0.1, we identified 9 network modules that gained connectivity (MDC > 1), but none of network modules lost connectivity (MDC < 1) under DR diet compared to control diet (Figure 4B. Supplementary File 2). Importantly, five of the differentially connected modules were also cycling modules identified from WGCNA, exhibiting higher network connectivity under DR (Figure 4B, Supplementary File 2). This suggests that DR may increase the diurnal coordination of gene expression in these modules.

**Figure 4.**
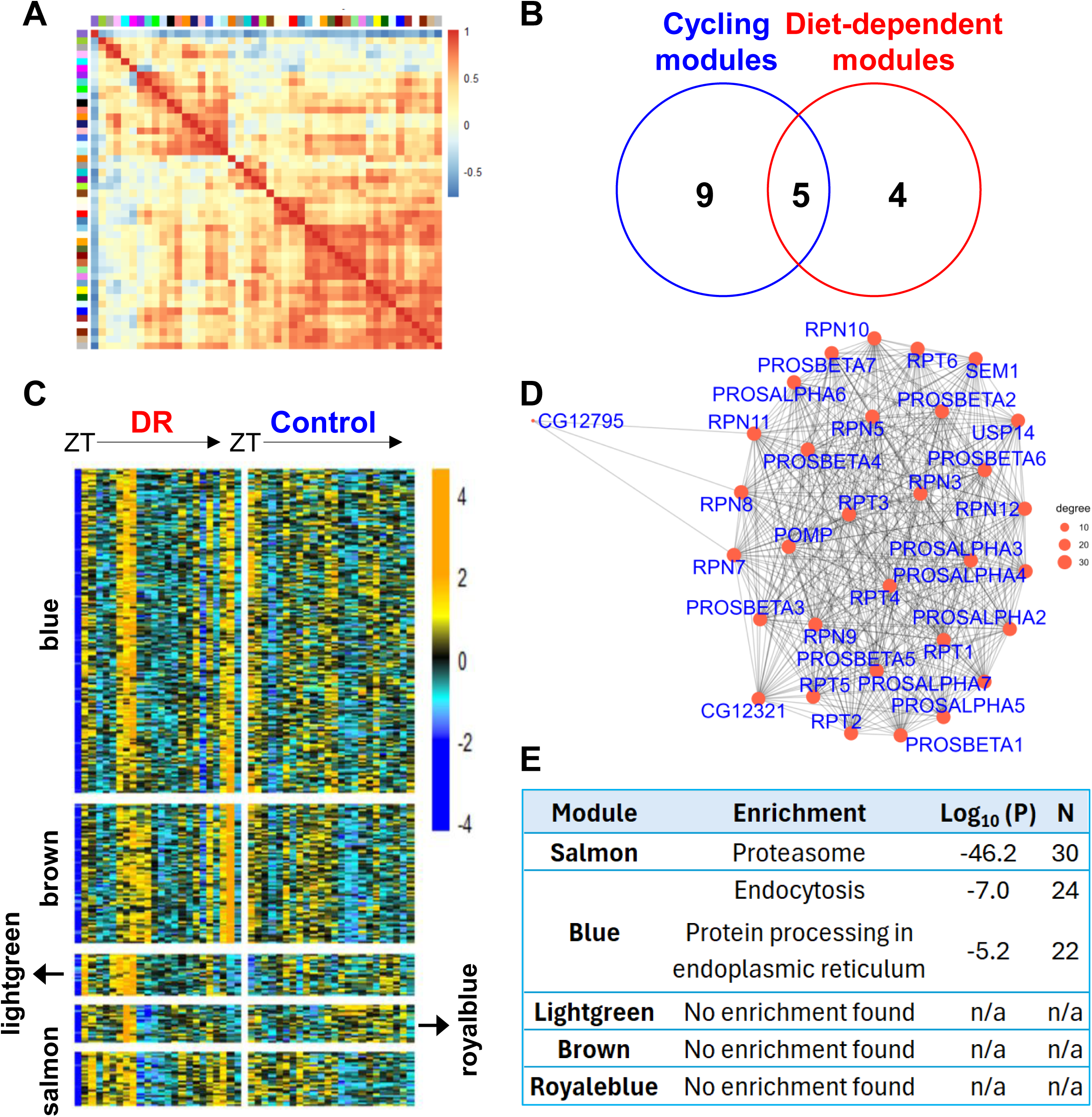
Identification of the “proteasome module” as a diet-dependent differentially regulated module enriched with oscillating genes. (A) Identification of a co-expression module enriched with proteasome subunit genes under DR. Gene co-expression networks under the DR diet were built using the WGCNA/r package. A topological overlap matrix (TOM) was then computed to evaluate the neighborhood similarity between genes and to classify network genes into modules using hierarchical clustering and dynamic tree cut (see Methods). (B) Venn Diagram shows the number of modules enriched with rhythmic genes (FDR < 0.05 and odd ratio > 0) and differentially regulated by diet (q < 0.1 and MDC > 1). Five modules were enriched with cycling genes and were deferentially connected between DR and control conditions. (C) Heatmap showing gene expression across two 24-h light/dark cycles in the five common modules. Each row represents a gene, and each column represents a sample ordered by sampling time. Expression levels were standardized across all samples and are presented as Z-scores. (D) Physical interaction map among the genes in the proteasome module in Bayesian network. Physical interaction mapping among the genes in the proteasome module was analyzed using a Bayesian network approach implemented in the bnlearn R package. (E) KEGG pathway enrichment analysis for the five differentially connected cycling modules. Enrichment scores in p-values were corrected using Benjamini-Hochberg approach with a threshold of 0.05.

One of these modules (the salmon module in Figure 4C), which we term the “proteasome module”, was of special interest. The protein products from many of the genes in this module physically interact with each other and constitute the core and auxiliary components of the proteasome complex, showing a dramatically higher pathway enrichment score than the other four modules (Figure 4D-E). A total of 30 of the 85 genes in this “proteasome” module were associated with the proteasome-mediated ubiquitin-dependent protein catabolic process (Figure 4D-E, Supplementary File 2). This observation also agrees with the primary analysis (Figure 3B, C) that, out of the 33 subunits of the proteasome complex in *Drosophila* (Belote and Zhong 2009), six proteasome subunits were defined as cycling (FDR < 0.1, log_2_ fold change > 0.6) on the DR diet while none of them cycled on control diet. When a less stringent rhythm detection cutoff (FDR < 0.2, log fold change > 0.6) was applied, the number of cycling genes under DR increased from 6 to 21, whereas none cycled under the control diet even with this more lenient threshold. Remarkably, 19 of these 21 genes were found in the proteasome module which gained network connectivity under DR, suggesting a DR-specific diurnal coordination. Thus, this observation suggests that circadian clocks modulate daily proteasome subunit gene expression in a nutrition-dependent manner.

### Knockdown of proteasome subunit genes in fat body limited normal lifespan and the effects of dietary restriction

Our finding of DR-induced changes in proteasome gene expression and cycling suggests a potential mechanism by which the clock could mediate DR effects. The proteasome complex functions as one of the major proteolytic degradation machines and is essential for maintaining cellular proteolysis (von Mikecz et al. 2008). Loss of protein homeostasis (proteostasis) is a hallmark of aging and can determine lifespan in both humans and model organisms, including flies (Kaushik and Cuervo 2015; Santra et al. 2019; Yang et al. 2019; Koyuncu et al. 2021; Meller and Shalgi 2021; Yu and Hyun 2021). The proteasome plays a central role in proteostasis by clearing, recycling, and breaking down up to ∼ 90% of cellular proteins (von Mikecz et al. 2008; Jang 2018). In multiple animal models, activation of the proteasome system extends lifespan and healthspan (Tonoki et al. 2009; Kruegel et al. 2011; Vilchez et al. 2012; Chondrogianni et al. 2015; Munkacsy et al. 2019; Anderson et al. 2022). For example, in *Drosophila*, global overexpression of *rpn11*, a regulatory subunit of the proteasome complex, extends lifespan while knock-down of *rpn11* decreases lifespan (Tonoki et al. 2009). Similarly, adult-specific ubiquitous overexpression of *pros*β*5*, a catalytic subunit of the proteasome, extends lifespan in flies (Nguyen et al. 2019). Moreover, pharmacological inhibition of the proteasome reduces lifespan in a dosage-dependent manner (Tsakiri et al. 2013). Intriguingly, DR extends lifespan in worms through promoting proteostasis, suggesting a link between DR and the proteasome (Matai et al. 2019). However, in flies, whether fat body proteasome function is linked to aging and/or the DR response has not been reported.

To gain insight into the role of the fat body proteasome, we used the mifepristone (RU486)-inducible GAL4/UAS Gene-Switch (GS) system (Osterwalder et al. 2001; Roman et al. 2001) to knockdown the expression of 11 proteasome subunits predominantly in the adult abdominal fat body using the S106-GS-Gal4 driver (Roman et al. 2001; Poirier et al. 2008; Bai et al. 2012; Jin et al. 2020; Taylor et al. 2022) and assess its effects on lifespan (Figure 5-figure supplement 1). Among those genes that displayed the most robust (>25% change) effects on lifespan, we focused on two genes, *pros*β*3* and *rpn7*, which are among the six genes that gained rhythmicity on the DR diet (FDR <0.1, log_2_ fold change > 0.6) While we observed some variability among independent trials (Figure 5-figure supplement 2), which has previously been observed in longevity assays in *Drosophila* (Bai et al. 2015; Katewa et al. 2016), we found significant reductions in lifespan as well as suppressions of DR-mediated lifespan extension for knockdown of both of these subunits (p < 0.0001 and p = 0.0222 for *pros*β*3* and *rpn7*, respectively for the gene×diet interaction effect from pooled data) (Figure 5, Figure 5-figure supplement 2, Supplementary File 1). In the case of *pros*β*3*, three out of the four trials demonstrated statistically significant gene×diet interactions (Figure 5-figure supplement 2, Supplementary File 1). Although the S106-GS-Gal4 is the most widely used inducible fat body driver, it also mis-expresses in the digestive system (Poirier et al. 2008). To test whether the reduced DR response by *pros*β*3* and *rpn7* knockdown with S106-GS-Gal4 is solely from the abdominal fat body or in the intestine or both, we performed two rounds (trial 1 and 2) of adult- and tissue-specific knockdown experiments using the gut-specific inducible TIGS-Gal4 (Poirier et al. 2008; Ulgherait et al. 2020). Although the DR effect was weaker in TIGS-Gal4 control flies (without RU486) compared to other wild-type control flies tested in this study (Figure 1, Figure 1-figure supplement 2, 4, Figure 5-figure supplement 1, 2), presumably due to genetic background effects (Liao et al. 2013; Jin et al. 2020; Wilson et al. 2020), knockdown of *pros*β*3* and *rpn7* in the gut consistently displayed a stronger lifespan extension by DR (Figure 5-figure supplement 3, Supplementary File 1) despite some variation in the lifespan pattern between the two trials. Notably, compared to S106-GS-Gal4 experiment (Figure 5, Figure 5-figure supplement 2), flies with either of the subunits knocked down in the gut suffered much stronger reductions in lifespan regardless of diet types (Figure 5-figure supplement 3), implying that the S106-GS-Gal4 results are not significantly affected by leaky misexpression in the gut. Thus, our data indicate that proteasome function in the adult abdominal fat body is important for DR-mediated lifespan extension.

**Figure 5.**
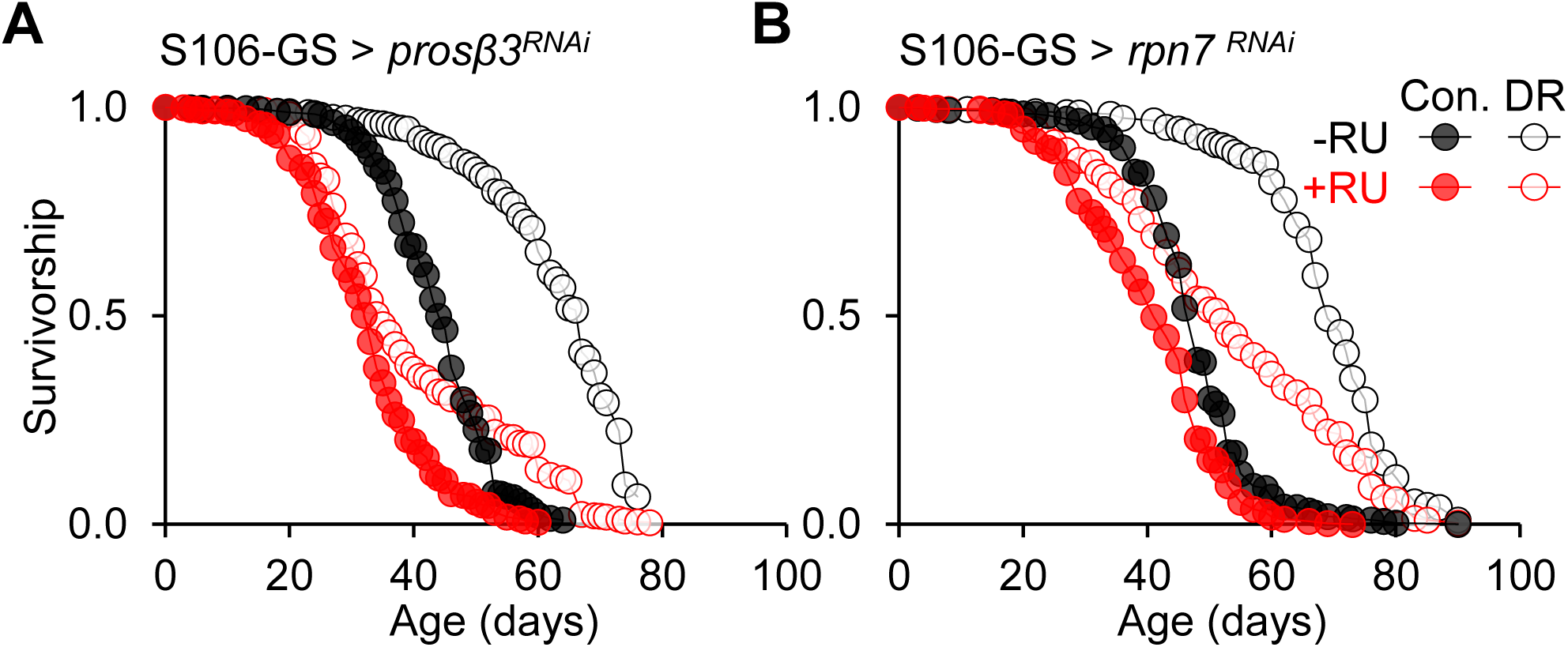
Reduced longevity response to DR by *pros*β*3* and *rpn7* knockdown in the abdominal fat body. (A–B) Survival curves of flies with *pros*β*3* and *rpn7* knockdown (+RU) in adult abdominal fat body (S106-GeneSwitch (GS) driver) and their controls (−RU) on control and DR diets. Survival curves were pooled from 3–4 independent trials (see also Figure 5-figure supplement 1, Supplementary File 1 for survival curves and detailed statistical analyses of independent trials). P < 0.0001 and p = 0.0169 for *pros*β*3* and *rpn7*, respectively, represent gene × diet interaction effects from Cox proportional hazards analysis.

## Discussion

Circadian rhythms dampen during aging, while environmental and genetic perturbations leading to high-amplitude circadian rhythms correlate with many health benefits (He et al. 2016; Froy 2018). Here we demonstrate that the master circadian clock transcription factor *Clk* is important for DR effects on lifespan and fecundity. We also discovered that DR acutely (∼ 5 days) alters the diurnal transcriptome in the fat body while sparing core clock genes, suggesting that the primary effects of DR are on rhythmic output genes. Using adult- and tissue-specific RNAi, we show that diet sensitive clock-controlled proteasome subunit genes in the abdominal fat body are important for the lifespan extending effects of DR. Our data suggest a role of the clock in DR effects on lifespan and reveal a molecular pathway, the proteasome, through which the clock may exert some of these effects.

Our data demonstrate a profound role for the transcription factor *Clk* in mediating the effects of DR on two independent phenotypes: lifespan and fecundity. The effects of *Clk^Jrk^*on DR-mediated lifespan expansion were robust, replicable, and, importantly, exhibited across a wide range of diets. These changes cannot simply be attributed to the shorter lifespan of *Clk^Jrk^* flies as *foxo* mutants also are short-lived but retain a robust DR-mediated lifespan extension (Giannakou et al. 2008; Min et al. 2008). Testing DR effects by using just two diets can mask a shift in the diet-dependent lifespan curve (Clancy et al. 2002; Flatt 2014), for example, due to changes in feeding. The fact that we observe a suppressed response to DR across a wide range of diets suggests a bona fide DR phenotype. In fact, *Clk* is one of just a handful of fly genes (Wang et al. 2009; Zid et al. 2009; Banerjee et al. 2012; Katewa et al. 2016; Ulgherait et al. 2016) for which this more rigorous standard has been achieved, suggesting a unique and central role of *Clk* in DR. *Clk^Jrk^* mutants are also not simply adversely affected by restricted nutrient intake, as may be the case for Bmal1 mutant mice (Kondratov et al. 2006), as *Clk^Jrk^* mutants are much more long-lived than *iso31* flies under the 1SY and 1Y malnutrition diets (Figure 1, Figure 1-figure supplement 2, 4).

Female fecundity in *Drosophila* is strongly correlated with diet concentration (Bass et al. 2007). *Clk^Jrk^* flies also exhibited a suppressed diet-dependent fecundity response especially at higher diet concentrations (Figure 2, Figure 2-figure supplement 1), resulting in a reduced fecundity response to a range of diets. These data suggest a more general role for *Clk* in dietary sensitivity beyond lifespan. Reduction of fecundity in female *Clk^Jrk^* mutants is consistent with previous observations under standard diets (Beaver et al. 2002).

While we observed reduced DR effects in *Clk^Jrk^* mutants, we observed a relatively robust DR lifespan response in arrhythmic *per^01^*mutants. Notably, our results reproduced one of the prior reports (Ulgherait et al. 2016) with *per^01^.* Loss of the major activator (*Clk*) and repressor (*per*) stop the clock at different stages of the cycle (Emery et al. 1998; Glossop et al. 1999; Claridge-Chang et al. 2001). As a result, *per* and *Clk* mutants can often yield distinct, even opposing, phenotypes (Keene et al. 2010). While we cannot exclude the possibility that *Clk^Jrk^* may not act via control of oscillatory gene expression (see (McDonald and Rosbash 2001)), we favor the idea that the clock drives daily oscillations between DR-sensitive (low PER) and DR-insensitive (low CLK) states which may be adapted to daily feeding (Xu et al. 2008) and/or the sleep/wake rhythm. Ultimately, circadian resonance experiments (e.g., (Xu et al. 2019)) are the gold standard to assess the role of circadian timing.

In addition to demonstrating a critical role for *Clk* in mediating the effects of DR on lifespan, we also find that DR reprograms the diurnal transcriptome, not by changing core clocks, but rather by inducing or altering rhythmicity of a key set of genes. In accordance with the established guidelines (Hughes et al. 2017), we collected samples every 2 h from two independent cycles of 24 h (every 2 h for 48 h), increasing the statistical power for rhythm detection, and computed the false discovery rate. To determine how the circadian clock may mediate DR effects, we assessed the circadian transcriptome under DR conditions in the fat body, a tissue important for metabolism, longevity, and DR. Importantly, we assessed the transcriptome after just 5 days of DR, sufficient time for DR to induce changes in mortality rate (Mair et al. 2003; McCracken et al. 2020). We found that this short-term DR was sufficient to produce 64% more rhythmic genes in total compared to control diet (Figure 3B). DR also increased the overall expression of many genes that are rhythmic under both conditions (Figure 3C, E-F). This is consistent with the observation in mouse liver that DR increases the number of rhythmic genes as well as their amplitude (Sato et al. 2017), indicating this feature of circadian DR sensitivity is widely conserved. Strikingly, although we discovered that DR for 5 days is sufficient to initiate reprogramming of the rhythmic transcriptome, core clocks remained virtually unaffected (Figure 3D), a time at which DR mortality effects are observed (Whitaker et al. 2014). In contrast, increased amplitudes of core clocks in mice liver is seen after 2∼ 6 months of DR (Patel et al. 2016b; Sato et al. 2017) and in flies after 10 days (Katewa et al. 2016).

Genetic induction of rhythmic amplitude of the core clock gene *timeless* altered lifespan in a DR-sensitive manner (Katewa et al. 2016). Yet, this *tim* induction was not accompanied by downstream changes in other components of the feedback loop, suggesting the core clock per se was not involved. Thus, we hypothesize that on the time scale of DR-induced changes in mortality rate, core clocks are not affected and that later changes in core clocks reflect an indirect and delayed effect of DR. It will be of interest to determine if short-term DR in mammals also spares core clock genes. Together, we postulate that core clocks are resistant to amplitude changes by short-term DR regimens while a longer term, depending on species, gradually increases their amplitude, which can further reprogram diet-dependent CCGs. This also implies that the duration of DR shapes the pattern of circadian transcriptome reprogramming.

Our data suggest a model by which the circadian clock gates the response to dietary restriction to control the complement and amplitude of clock regulated gene expression (Figure 6). First, the circadian clock rhythmically controls the daily expression of multiple components of the proteasome. Second, DR increases the expression levels and enhances coordinated rhythmic expression of proteasome subunit genes. WGCNA followed by MDC analysis discovered that the transcriptomic organization in the proteasome module was altered by DR. This change is due to enhanced and coordinated rhythmic expression of proteasome subunits by DR (Figure 4). Importantly, we provide in vivo evidence that these subunits of the proteasome in the fat body are important for diet sensitive effects on lifespan, providing a pathway of linking clocks, DR, and aging.

**Figure 6.**
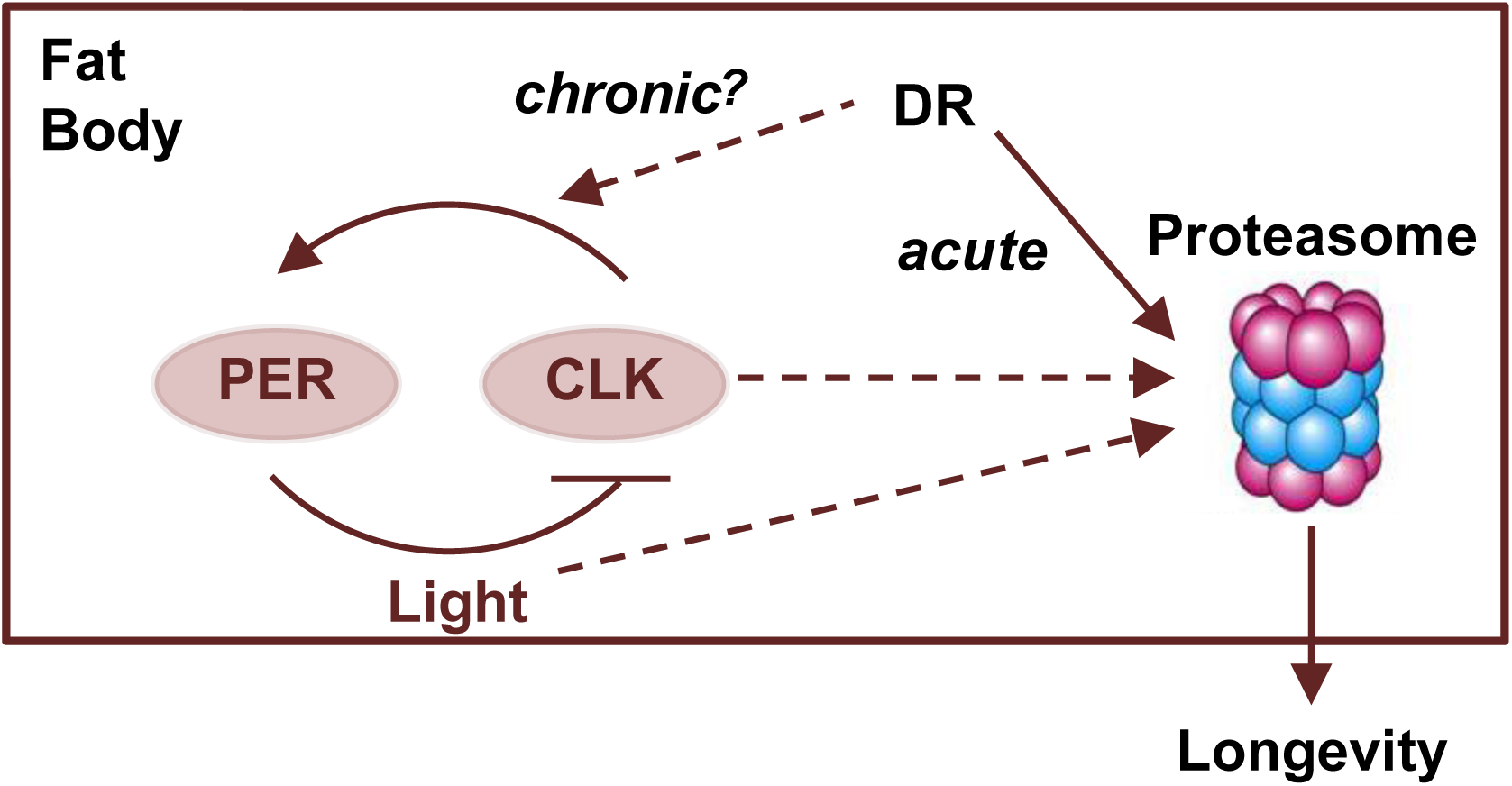
**Model for how clock and diet impact proteasome expression to regulate lifespan** Acute DR increases the number of diurnally rhythmic genes, including proteasome genes, without significantly affecting core clock genes in the abdominal fat body. The diet-dependent gain of oscillation in proteasome genes enhances proteostasis and contributes to DR-mediated lifespan extension. In contrast, prolonged DR may further modulate the circadian clock itself, potentially increasing the rhythmicity of clock-output genes.

Although it is well-established in multiple species that proteasome activity decreases during aging and is generally positively correlated with lifespan (Tonoki et al. 2009; Kruegel et al. 2011; Vilchez et al. 2012; Chondrogianni et al. 2015; Pickering et al. 2015; Huang et al. 2019; Nguyen et al. 2019; Anderson et al. 2022), little is known about its tissue-specific contribution to aging and the link to DR. We found that the knock-down of several cycling proteasome subunits predominantly in the adult fat body, using the S106-GS Gal4 (Poirier et al. 2008) (Figure 5-figure supplement 1), significantly reduces lifespan, consistent with whole organism manipulations (Tonoki et al. 2009). Moreover, further testing for potential DR effects with two selected subunits (*pros*β*3* and *rpn7*) revealed that knock-down of these subunits reduced DR effects, providing evidence that suppression of proteasome function in the fat body limits DR-mediated lifespan extension (Figure 5, Figure 5-figure supplement 2). While some variability was observed, perhaps due to inconsistent delivery of RU486 to flies (Yamada et al. 2017), in the case of *pros*β*3*, significant effects on DR response were observed in three out of four trials (Figure 5-figure supplement 2). Moreover, combined data from independent trials confirmed knockdown of these subunits reduces the DR effect (Figure 5). It will be of interest if these effects on DR-sensitive longevity also extend to fecundity. We favor the idea that impaired proteasome function contributes to amino acid imbalance, which is known to be critical for DR-mediated lifespan extension in flies (Grandison et al. 2009), leading to reduced DR response (Figure 6).

As DR is suggested to be the most promising intervention to delay aging, the work presented here has important implications for integrating timing into anti-aging therapies. In conjunction with our observations, the beneficial effects of time-restricted feeding on longevity is strongly abolished in core clock mutant flies including *Clk^Jrk^* flies (Ulgherait et al. 2021), emphasizing the roles of circadian clocks in dietary interventions for health and longevity. We propose that time-of-day activation of proteasomes may, at least partially, mediate the beneficial effect of DR. Thus, the daily timing of anti-aging therapies may be crucial for lifespan and healthspan extension.

## Materials and Methods

### Fly Rearing and Media

All the flies used for experiments were raised on a standard yeast-cornmeal-molasses based diet (per 1 L water: agar 12.9 g, soy flour 22.3 g, yeast 38.6 g, cornmeal 162.9 g, molasses 85.7 mL, corn syrup 85.7 mL, nipagin (methylparaben) 2.9 g, ethanol 26.8 mL, and propionic acid 10.7 mL) under a light-dark (LD) 12:12h cycle. The following flies were used in this study. *Clk^Jrk^* and *per^01^* flies were backcrossed to the wild type (*w^1118^*) *iso31* line (BDSC# 5905) 6 times. S106-GeneSwitch (S106-GS) (BDSC# 8151) and RNAi lines were obtained from Bloomington Stock Center (Supplementary File 1). The TIGS-Gal4 driver line was generously provided by Dr. Hua Bai (Iowa State University).

### Lifespan and Fecundity Assay

For the lifespan assay, young adult female flies (∼ 48 hours cohorts) were separated under light CO_2_ anesthesia after mating with males for ∼ 2 days in food bottles. Separated female flies were kept in groups of 20∼25 flies in plastic vials on the sucrose-yeast (SY) diet (per 1 L: agar (Genesee Scientific # 66-103) 15 g, sucrose and yeast (Brewers yeast, MP Biomedical) added at a 1:1 ratio ranging from 10–200 g each depending on diet concentration, ethanol 15 mL, propionic acid 10 mL, and nipagin 3 g) and transferred to fresh food vials every 2∼3 days. For the lifespan experiment with Mifepristone (RU486)-inducible GeneSwitch system, flies were kept in the vials containing either vehicle (1% EtOH) or RU486 (200 uM). RU486 or EtOH was thoroughly mixed into the experimental food before it was dispensed into vials. Dead flies were recorded at each transfer. For the fecundity assay, single females (∼ 3 days old) were placed with two males of similar age in vials containing different SY diets. Vials were changed daily at ZT0 (8 AM) for 7 days and stored at 4°C until the eggs were counted. Vials with dead females or sterile females (no eggs laid) during the assay were removed from the analysis. For both lifespan and fecundity assays, flies were kept at 25°C, 12hr light : 12 dark (12L:12D) and 60% relative humidity.

### Survival Statistics

Survival analysis, log-rank test to evaluate statistical differences between survival curves, and Cox proportional hazards analysis to evaluate ability of tested genes to modify lifespan in the specified range of diets were performed with JMP® statistical package (version 14, SAS Institute Inc.) with data from replicate vials combined. P values from the Cox proportional hazards analysis represent the probability of main (Gene (G) as a nominal variable and Diet (D) as a continuous variable) and interaction effects (G x D) by the likelihood ratio chi-square test.

### Fat Body Dissection

Young mated female flies (∼ 3 day old) were entrained under either DR diet (5SY) or control diet (15SY) for 5 days in 12L:12D cycles at 25°C. At every 2 hours, flies were directly dissected without dry ice to harvest fat tissues in the abdomen. Pinned flies were cut to remove organs in the abdomen (intestine, ovaries, Malpighian tubules, etc). Fat tissues attached to the epidermis were collected (Xu et al. 2008; DiAngelo and Birnbaum 2009; Xu et al. 2011; Katewa et al. 2016). Fat body from ∼ 10 flies were harvested within 10 minutes for each time point of RNA-Seq analysis.

### RNA-Seq

Dissected fat body tissues from ∼8 days old mated female flies were homogenized in pH 7.4 PBS for 2 min using a Kontes motor and pestle, and were incubated TRIzol LS Reagent (Thermo Fisher, Waltham, MA) for 15 min. RNA was extracted according to the manufacturer’s instructions and residual DNA in the homogenized samples was removed by RNAse free DNase I (Thermo Fisher Scientific). Quality of RNA samples were checked with Agilent 2100 Bioanalyzer. cDNA library was constructed with poly(A) selected mRNA using Truseq RNA library preparation kit and then sequenced at the Genomics Core Facility at the University of Chicago on Illumina HiSeq 2000 System.

### Quantification of Transcript, Normalization, and Batch Correction

Raw reads were pseudo-aligned and quantified using Kallisto (v0.50.1) against a prebuilt index generated from the Ensembl reference transcriptome (release v96). For quantification, single-end reads were processed with 50 bootstrap replicates and an estimated average fragment length of 200 bp. Gene-level abundances were computed by summing transcripts per million (TPM) values across all transcripts belonging to a given gene. Genes with <1 TPM in more than 40% of time points per condition (including replicates) were excluded from further analyses. Batch correction was performed at the count level (est_counts), and corrected TPM values were subsequently estimated from these batch-adjusted counts. Normalization within each condition was performed using the RUVSeq package (Risso et al. 2014) under the RUVg protocol, employing a set of “in silico empirical” negative control genes (i.e., the least significantly differentially expressed genes identified in a first-pass DE analysis). Technical batch effects across conditions were adjusted using the EDASeq protocol (Risso et al. 2011) with upper-quartile (UQ) normalization.

Final corrected TPM values were calculated as:

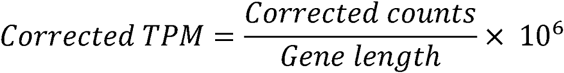

### Rhythmicity Detection

Rhythm detection was performed using RAIN (Thaben and Westermark 2014) on filtered TPM level data. Parameters were set for detection of a 24-h period, with a sampling interval of 2 h for the DR and control datasets and 4 h for the Clk^Jrk^ dataset. Genes with FDR corrected p-value < 0.1 and 0.6 > log2 fold change in raw TPM were assumed to be cycling.

### WGCNA and MDC

We reconstructed gene coexpression networks under the DR condition using the WGCNA/r package (Zhang and Horvath 2005; Langfelder and Horvath 2008). Briefly, we first computed the network adjacency matrix as kij = [0.5 * (1 – rij)] ^ β. kij is the network connectivity and rij is the Pearson correlation coefficient between a pair of genes i and j. Soft power threshold β was chosen so that the topology of the network was scale-free. A topological overlap matrix (TOM) was then computed to evaluate the neighborhood similarities between genes and to classify network genes into modules using hierarchical clustering and dynamic tree cut. We used DAVID (v24.1) to functionally annotate the identified network modules. To identify network modules that were organized by circadian rhythmicity, we tested the enrichment of rhythmically expressed genes (FDR < 0.1 identified using concatenated data from both DR and control diets) in network modules using fisher’s exact test. Since we defined the network as a “signed” network (i.e., using kij = [0.5 * (1 – rij)] ^ β, instead of the default choice of kij = |rij| ^ β), genes in the cycling network modules shared highly similar circadian phases (as opposed to also including genes with the exact opposite phases) as well as cycling waveforms. To evaluate changes in network connectivity between DR and control diets, we implemented in R a method to compute modular differential connectivity (MDC) described by Zhang et al., 2013 (Zhang et al. 2013). MDC is defined as the ratio of the summed pairwise connectivity among genes in a module under DR conditions and that of the same genes under control conditions (i.e., MDC = ∑i ∑j kijDR / ∑i ∑j kijcontrol). Statistical significance was determined using a permutation-based FDR approach. Two types of FDR estimates were computed, one based on randomly permuted samples to generate networks with nonrandom nodes but random connections, and the other based on randomly permuted gene labels to generate networks with random nodes but nonrandom connections. The final FDR was determined as the larger of the two estimates. 1000 permutations were computed for each type of FDR estimates, and FDRs for modules that gained connectivity (MDC > 1) or lost connectivity (MDC < 1) under DR conditions were estimated separately as described by Zhang et.al. (Zhang et al. 2013).

### Differential Expression Analysis

Differential gene expression analysis was performed using DESeq2 library (Love et al. 2014). We used est. count data generated by kallisto, and a union of pre-filtered gene lists for both conditions (gene lists were in agreement with the TPM level pre-filtering, as described above) resulting in 8440 gene IDs. The full model included time, in a form of 3rd degree polynomial, and diet type as factors.

### Data and Code Availability

The transcriptomic datasets used in this study are publicly available. Raw sequencing data (FASTQ files) for wild-type flies under control and dietary restriction conditions are deposited in the Gene Expression Omnibus under accession number GSE145509. Raw data for ClkJrk mutant flies are available under accession number GSE241003. Processed data, including TPM and estimated count matrices, as well as all scripts used for data processing and anCalysis, are available on GitHub at: https://github.com/shijusisobhan/DR-PeripheralClock-Longevity

## Acknowledgments

We thank Besim Becoja, Eric Ho Cheung, Alejandra Diaz, Gwang Min Han, Alexandra Raymond, Benjamin Green, Tova Beeber, and Mubaraq Opoola for technical assistance for lifespan and feeding assays and the Bloomington Stock Center for RNAi lines. This work was supported by Defense Advanced Research Projects Agency (DARPA) (D12AP00023 to RA), NIH R35 NS132223 to RA and the Data Science Initiative (DSI) Research Support Program (to RA) at Northwestern University. This effort was in part sponsored by DARPA; the content of the information does not necessarily reflect the position or the policy of the government, and no official endorsement should be inferred. This work was supported by the NSF-Simons Center for Quantitative Biology at Northwestern University. This work was supported by a grant from the Simons Foundation (597491-RWC) and the National Science Foundation (1764421). The content is solely the responsibility of the authors and does not necessarily represent the official views of the National Science Foundation and Simons Foundation. DH was supported by the National Institutes of Health T32 Institutional Training Grant (Northwestern Univ: grant NIH T32HL007909), the National Institute of General Medical Sciences of the National Institutes of Health under award number P20GM103436 (Univ. of Louisville), the National Institute on Aging under award number R15AG074538 (to DH). ALH was supported by the National Institutes of Health Medical Scientist Training program at the University of Chicago (grant NIGMS T32GM07281). Stocks obtained from the Bloomington Drosophila Stock Center (BDSC) (NIH P40OD018537) were used in this study.

## Supplementary Files

**Supplementary File 1. Summary of Descriptive Statistical Analysis of Lifespan Data. Supplementary File 2. Summary of Weighted Gene Co-expression Network Analysis (WGCNA) and Module Differential Connectivity (MDC).**

---

**Figure 1-figure supplement 1.**
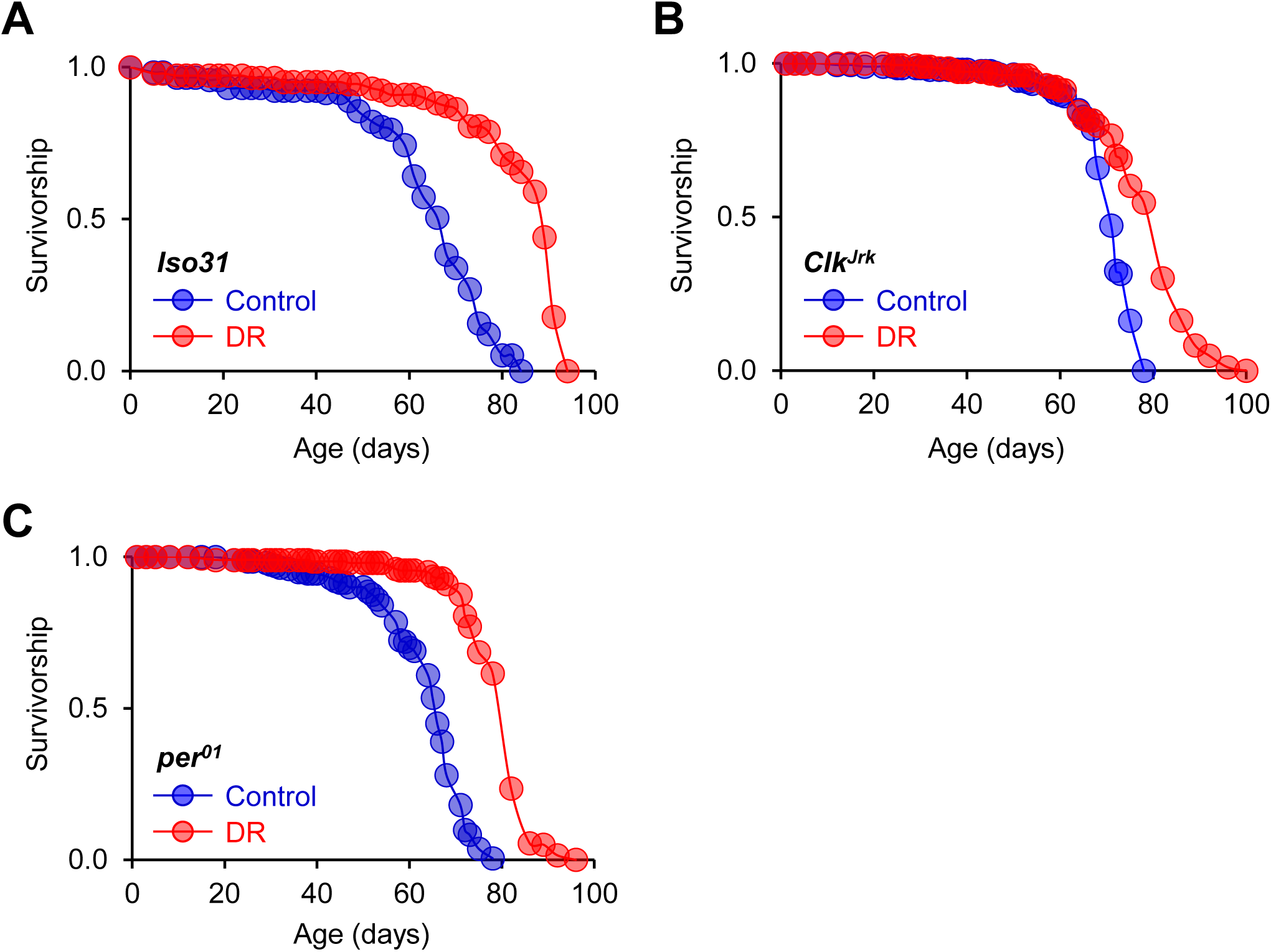
Reduced DR effect in *Clk^Jrk^* mutant flies compared to *iso31* wild-type and *per^01^* mutant flies. (A-C) Survival curves of *iso31*, *Clk^Jrk^* and *per^01^*homozygous mutant flies on control (15% sucrose-yeast; 15SY) or DR (5% sucrose-east; 5SY) diets. DR extended lifespan in *iso31*, *per^01^*, and *Clk^Jrk^* by 29.8%, 24.8%, and 11.3%, respectively. *n* = 117–200 flies (Supplementary File 1). Statistical significance of the diet x genotype interaction for ability to extend lifespan by DR between genotypes was analyzed by Cox proportional hazards regression analysis. p < 0.0001 for *iso31* vs *Clk^Jrk^* and *Clk^Jrk^*vs *per^01^*, and p = 0.0635 for *iso31* vs *per^01^*. See Supplementary File 1 for additional details of the statistical analysis.

**Figure 1-figure supplement 2.**
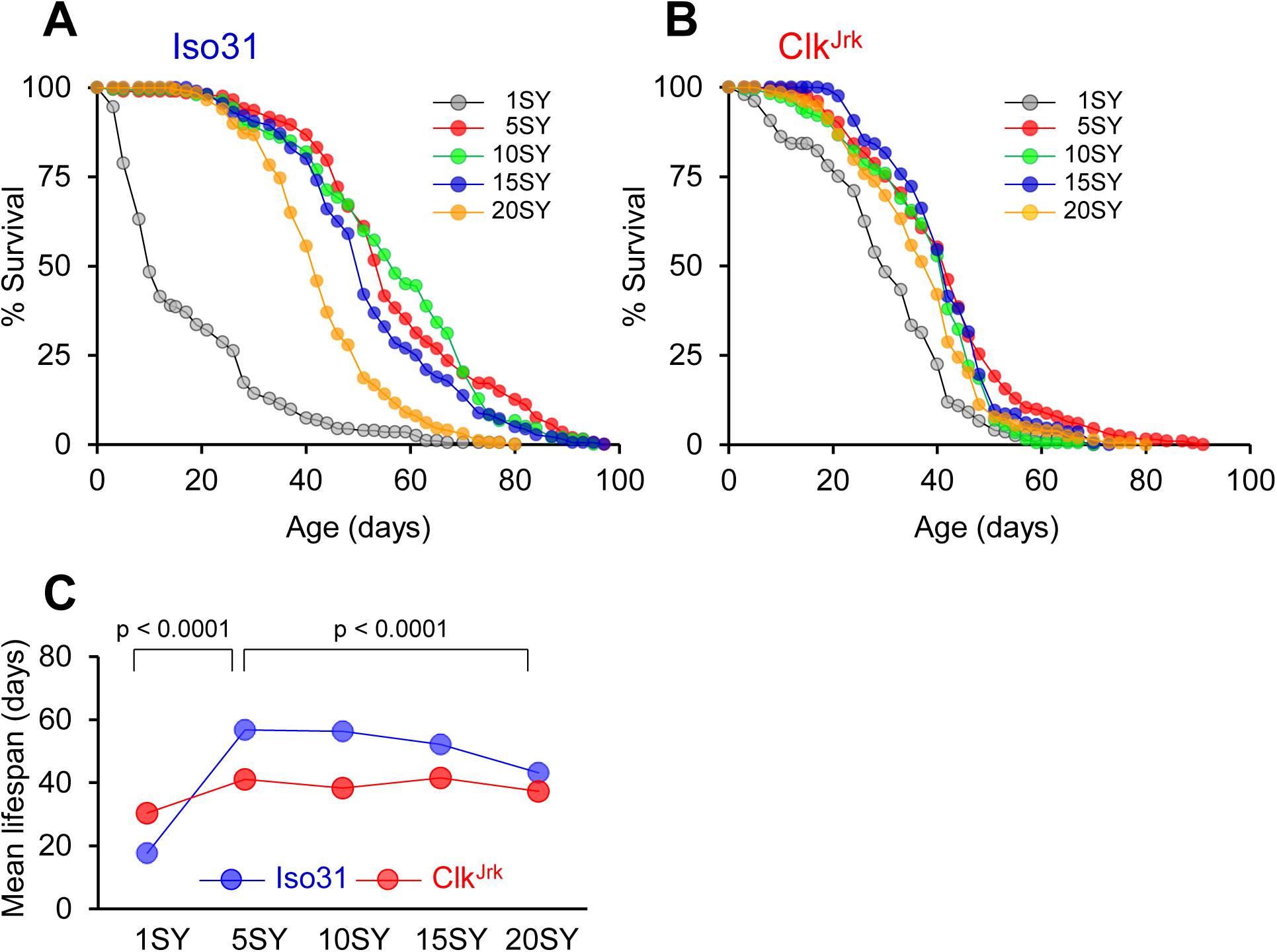
Reduced longevity response to DR in *Clk^Jrk^* mutant flies. Independent replication of the experiment presented in Figure 1. (A and B) Survival curves of wild-type control flies (*iso31*) and *Clk^Jrk^* homozygous mutant flies on 1%, 5%, 10%, 15%, and 20% sucrose-yeast (SY) diets. *n* = 198–208 flies (Supplementary File 1). (C) Mean lifespan plots of *iso31* and *Clk^Jrk^* flies across different concentrations of SY diets. P values represent the probability of main (Gene (G) as a nominal variable and Diet (D) as a continuous variable) and interaction effects (G x D) by the likelihood ratio chi-square test from Cox proportional hazards regression analysis to evaluate ability of tested genes to modify lifespan in the specified range of diets. Separate analyses were performed for normal DR response (from 5SY to 20SY) and malnutrition-like response (from 5SY to 1SY). See Supplementary File 1 for additional details of the statistical analysis.

**Figure 1-figure supplement 3.**
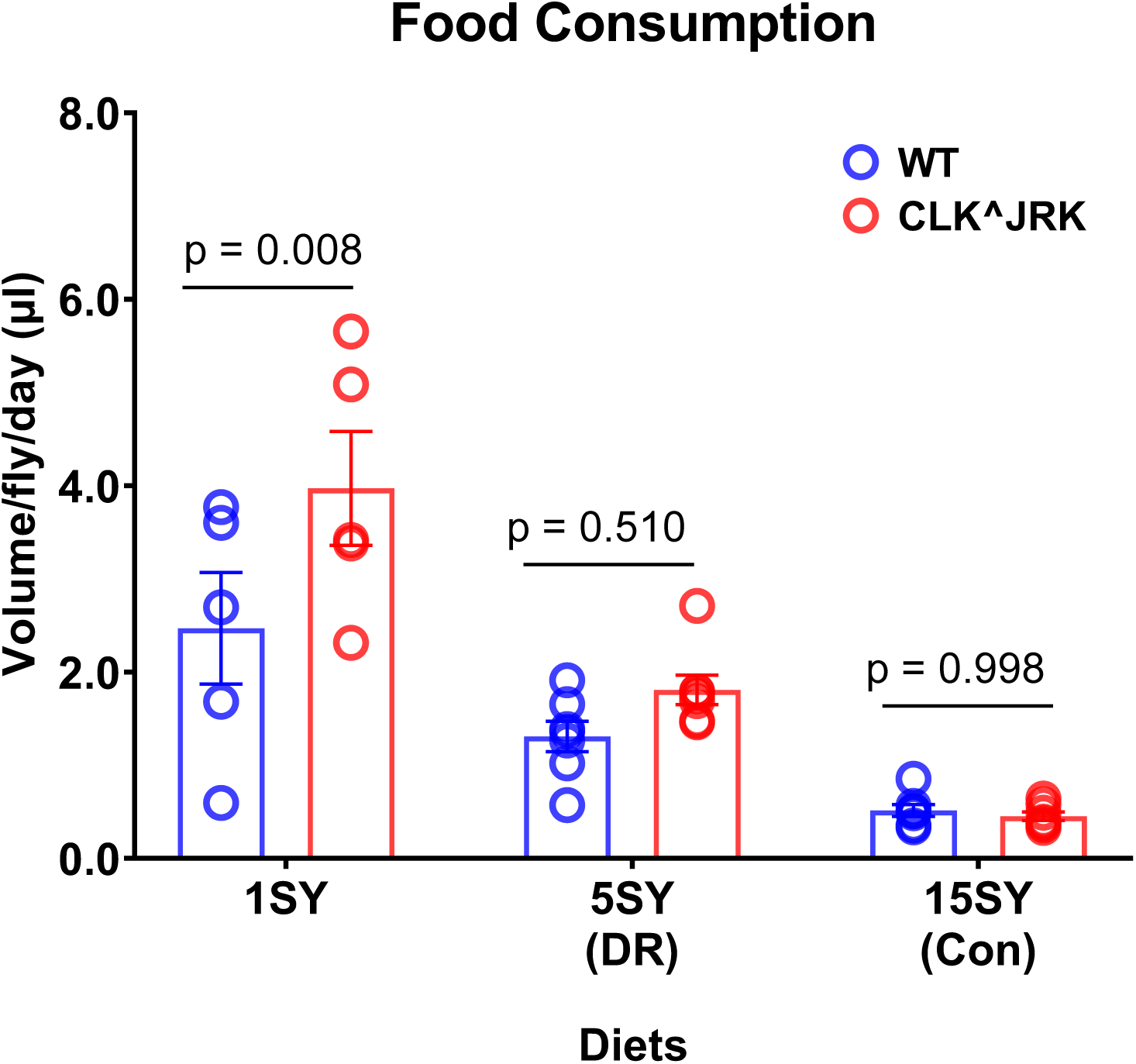
**Food Consumption in wild-type (*iso31*) and *Clk^Jrk^* flies** Young mated females (∼2 days old) were maintained on a designated diet for 5 days and then transferred to the same diet containing 1% blue dye (FD&C Blue No. 1) for 48 hours. The amount of dye in both internal tissues and excreta was quantified using a spectrophotometer at 630 nm and calculated using a standard curve. n = 5 replicates (3 flies per replicate). Two-way ANOVA with Sidak’s multiple comparisons: Diet: p < 0.0001, Genotype: p = 0.0112, Diet × Genotype interaction: p = 0.0476. A significant difference between WT and Clk^Jrk^ flies was observed only in the 1SY malnutrition diet, but not in the DR or control diets.

**Figure 1-figure supplement 4.**
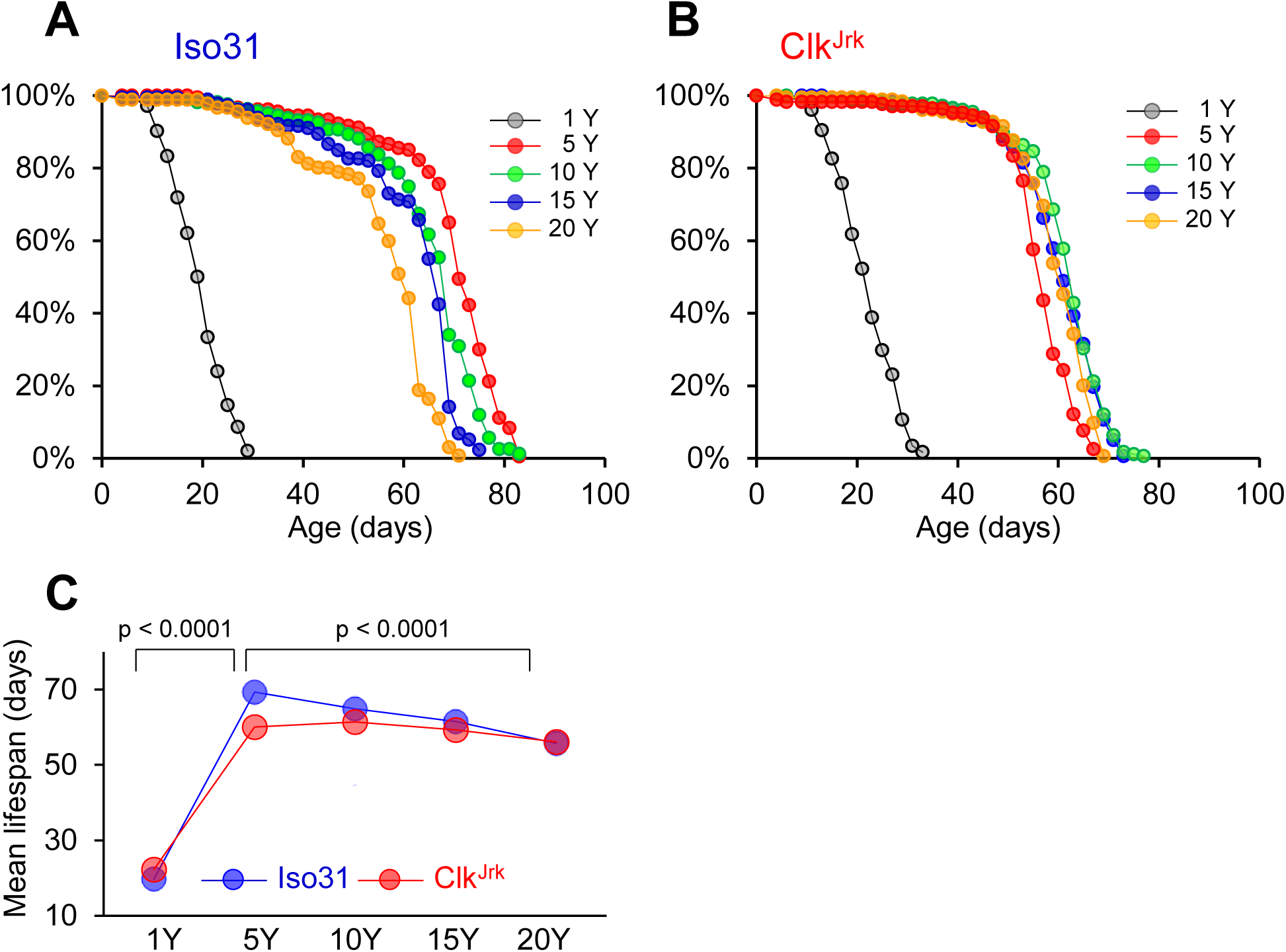
**Reduced longevity response to yeast (Brewer’s yeast; whole cell lysates) restriction in *Clk^Jrk^* mutant flies.** (A and B) Survival curves of wild-type control flies (*iso31*) and *Clk^Jrk^*homozygous mutant flies on 1%, 5%, 10%, 15%, and 20% yeast-restricted (Y) diets with a fixed sucrose concentration of 5%. *n* = 175–181 flies (Supplementary File 1). (C) Mean lifespan plots of *iso31* and *Clk^Jrk^*flies across different concentrations of Y diets. P values represent the probability of main (Gene (G) as a nominal variable and Diet (D) as a continuous variable) and interaction effects (G x D) by the likelihood ratio chi-square test from Cox proportional hazards regression analysis to evaluate ability of tested genes to modify lifespan in the specified range of diets. Separate analyses were performed for normal DR response (from 5Y to 20Y) and malnutrition-like response (from 5Y to 1Y). See Supplementary File 1 for additional details of the statistical analysis.

**Figure 1-figure supplement 5.**
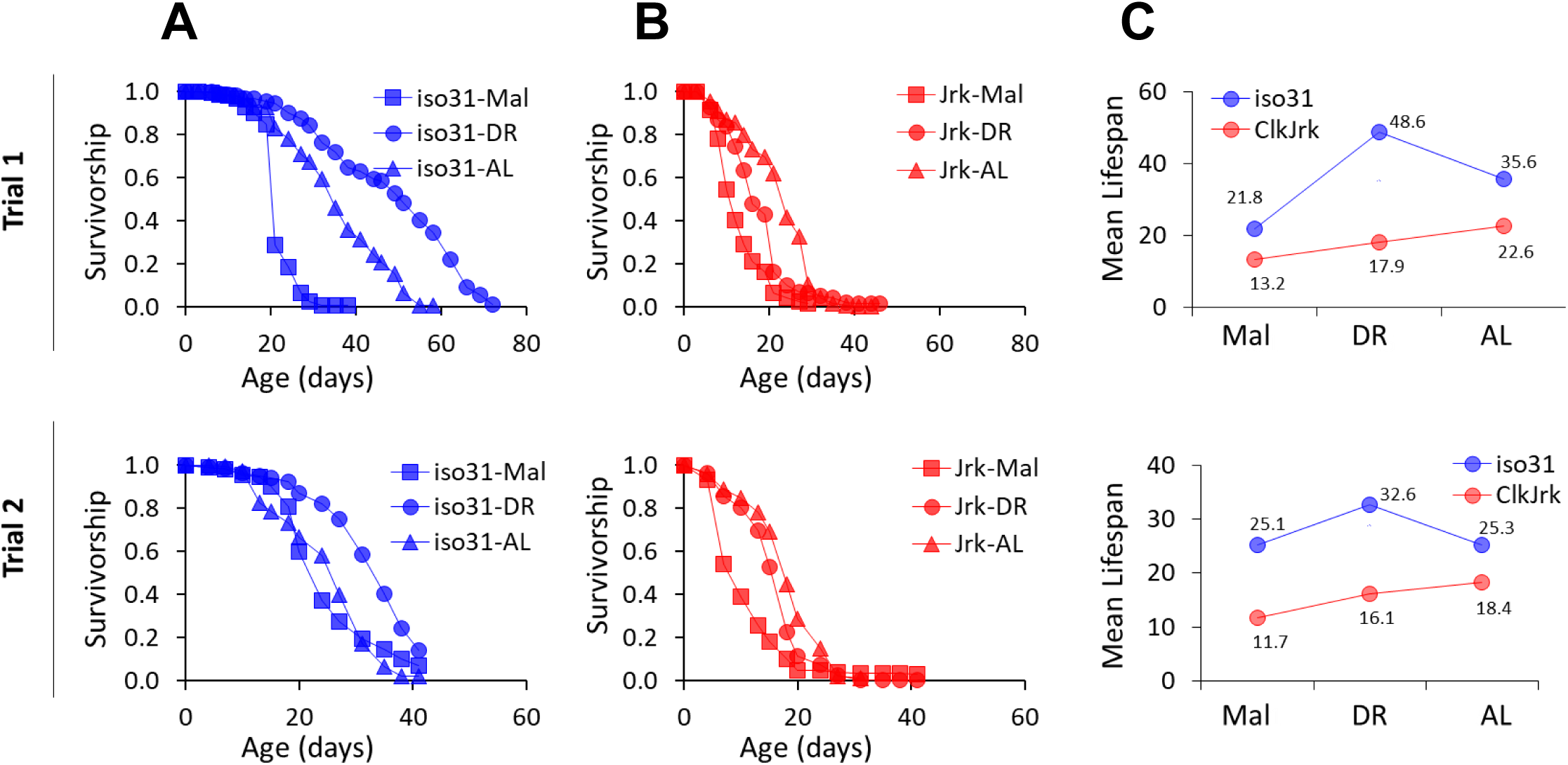
**Reduced longevity response to yeast (yeast extract) restriction in *Clk^Jrk^*mutant flies.** (A and B) Survival curves of wild-type control flies (*iso31*) and *Clk^Jrk^*homozygous mutant flies in varying concentrations of yeast extract with a fixed sucrose concentration of 5%. (C) Mean lifespan plots of *iso31* and *Clk^Jrk^*(Jrk) flies across different concentrations of yeast extract diets. Mal (malnutrition): 0.01%, DR: 0.5%, AL (ad libitum): 5% yeast extract. Upper and lower panels are from independent trials 1 and 2. See Supplementary File 1 for additional details of the statistical analysis.

**Figure 2-figure supplement 1.**
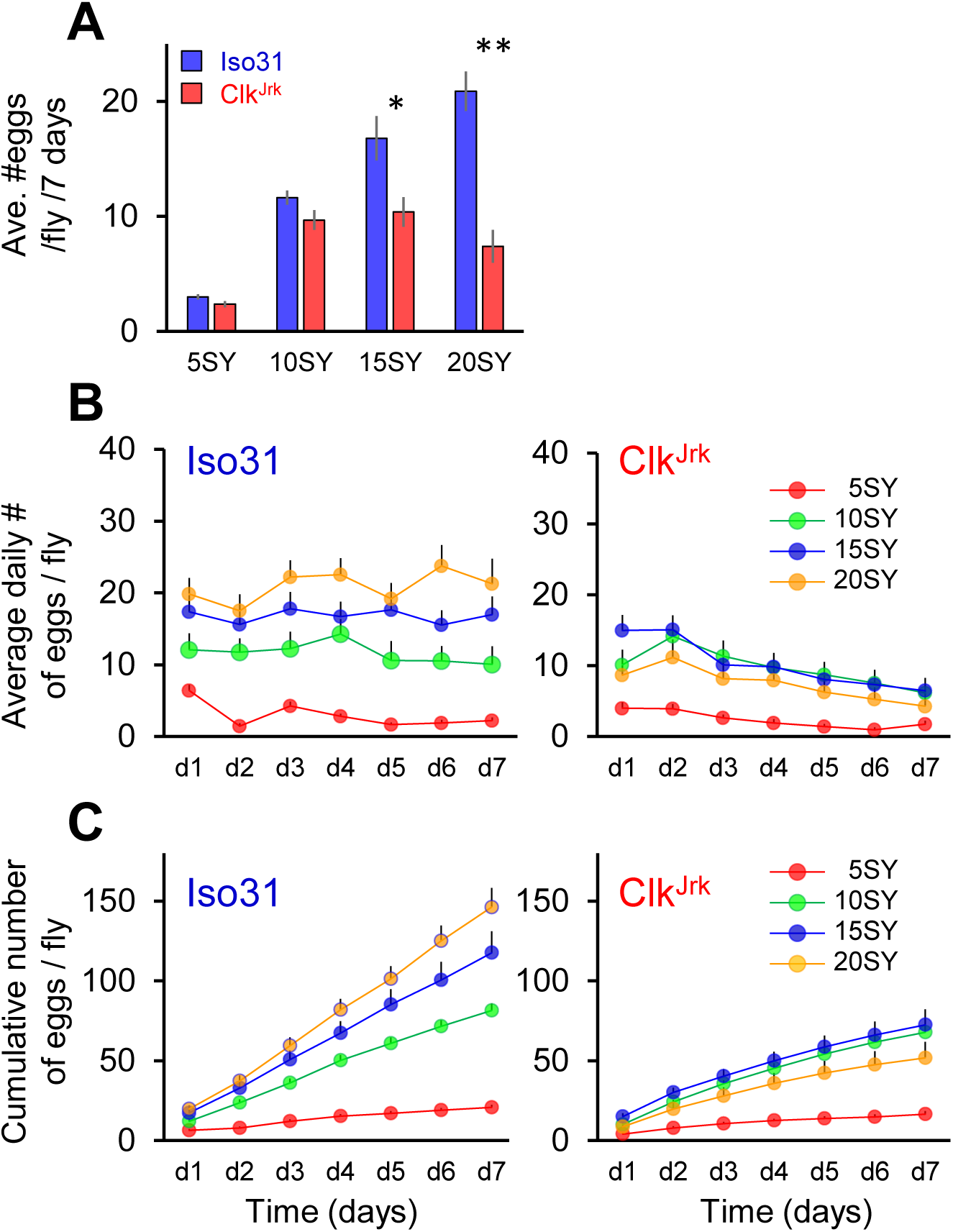
Reduced fecundity response to diets in *Clk^Jrk^* mutant flies. Independent replication of the experiment presented in Figure 2. (A and B) Cumulative and daily average number of eggs produced per fly over 7 days in wild-type control flies (*iso31*) and *Clk^Jrk^* homozygous mutant flies on 5% (n = 17, *iso31*; n = 18, *Clk^Jrk^*), 10% (n = 19, *iso31*; n = 19, *Clk^Jrk^*), 15% (n = 19, *iso31*; n = 13, *Clk^Jrk^*), and 20% (n = 16, *iso31*; n = 18, *Clk^Jrk^*) sucrose-yeast (SY) diets. (C) Cumulative number of eggs produced per fly over 7 days. * p < 0.05, ** p < 0.01 by t-test for specified pair-wise comparisons.

**Figure 3-figure supplement 1.**
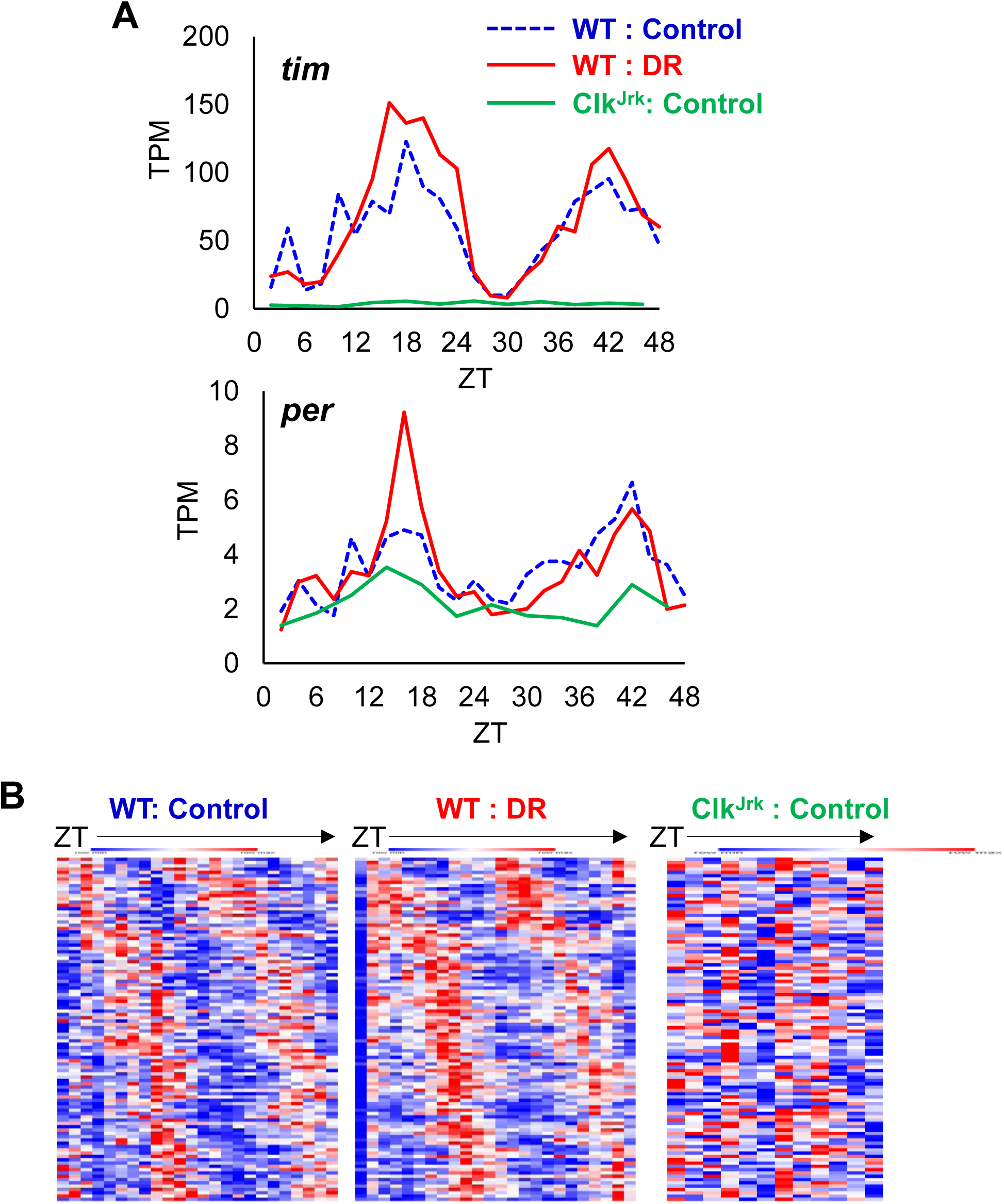
**Clock is required for diurnal rhythmic gene expression in the abdominal fat body.** (A) Diurnal expression patterns of core clock genes timeless (tim) and period (per) in WT and Clk^Jrk^ flies. (B) Diurnal rhythmic expression patterns (red = high, blue = low) of common cyclers in WT flies under control and DR diets, and their expression pattern in Clk^Jrk^ mutants under control diet. The heatmap displays relative expressions over 48 hours (2-hour intervals for WT; 4-hour intervals for Clk^Jrk^).

**Figure 5-figure supplement 1.**
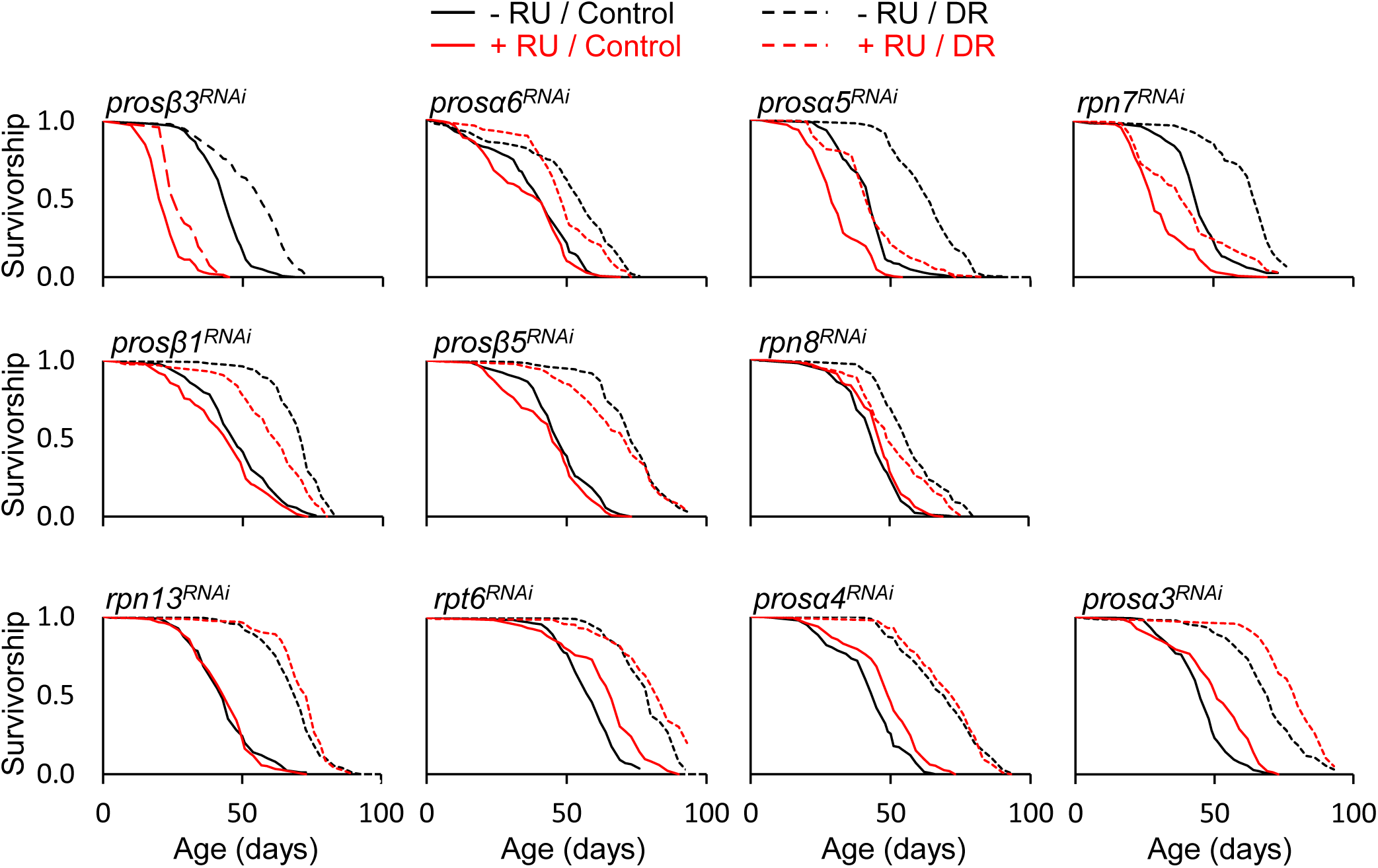
**Effect of proteasome suppression in the abdominal fat body on lifespan and DR.** Proteasome subunits were individually knocked down in the adult abdominal fat body with the S106-Gene Switch (GS) driver. Mated females were aged on either control (15% sucrose-yeast) or DR (5% sucrose-yeast) diets and provided fresh food every 2∼3 days (-RU: 1% EtOH, +RU: 200 uM RU486 in 1% EtOH). *n* = 104–185. See Supplementary File 1 for additional details of the statistical analysis.

**Figure 5-figure supplement 2.**
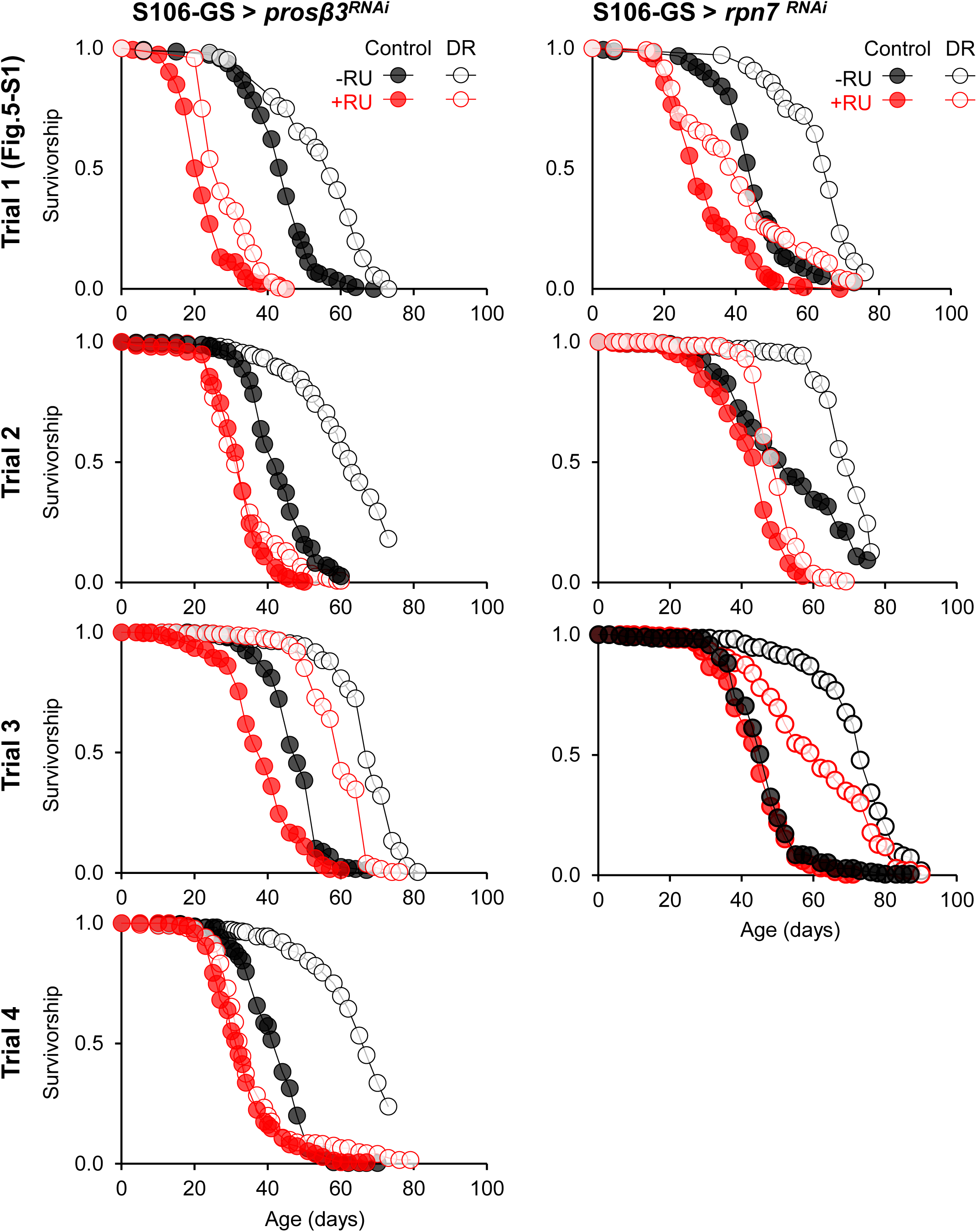
**Effect of *pros*β*3* and *rpn7* knockdown in the abdominal fat body on lifespan and DR.** Proteasome subunits (*pros*β*3* and *rpn7*) were individually knocked down in the adult abdominal fat body using the S106-GS driver. Mated females were aged on either control (15% sucrose-yeast) or DR (5% sucrose-yeast) diets and provided fresh food every 2∼3 days (-RU: 1% EtOH, +RU: 200 uM RU486 in 1% EtOH). For comparison among independent trials, the graphs for *pros*β*3* and *rpn7* from Figure 5-figure supplement 1 are presented again as trial 1. *n* = 104–294 in independent trials. See Supplementary File 1 for additional details of the statistical analysis.

**Figure 5-figure supplement 3.**
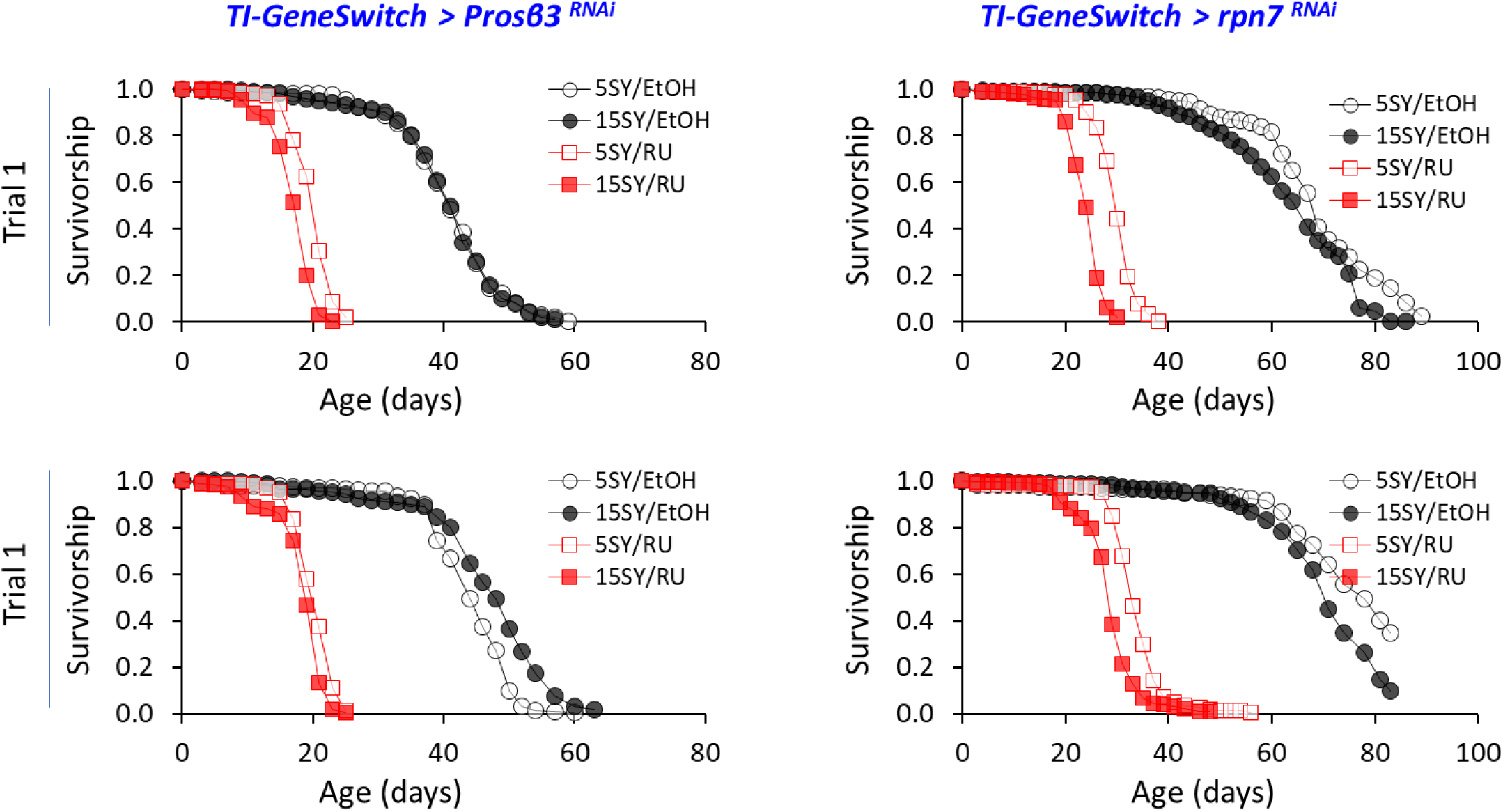
Effect of prosβ3 and rpn7 knockdown in the gut on lifespan and DR. Proteasome subunits (prosβ3 and rpn7) were individually knocked down in the adult gut using the TI-GS driver. Mated females were aged on either control (15% sucrose-yeast) or DR (5% sucrose-yeast) diets and provided fresh food every 2∼3 days (-RU: 1% EtOH, +RU: 200 uM RU486 in 1% EtOH). n = 223–253. See Supplementary File 1 for additional details of the statistical analysis.

